# Phylogenetic Clustering by Linear Integer Programming (PhyCLIP)

**DOI:** 10.1101/446716

**Authors:** Alvin X. Han, Edyth Parker, Frits Scholer, Sebastian Maurer-Stroh, Colin A. Russell

**Author notes:** These authors contributed equally to this work (Alvin X. Han, Edyth Parker). Corresponding authors: Alvin X. Han and Colin A. Russell.

## Abstract

Sub-species nomenclature systems of pathogens are increasingly based on sequence data. The use of phylogenetics to identify and differentiate between clusters of genetically similar pathogens is particularly prevalent in virology from the nomenclature of human papillomaviruses to highly pathogenic avian influenza (HPAI) H5Nx viruses. These nomenclature systems rely on absolute genetic distance thresholds to define the maximum genetic divergence tolerated between viruses designated as closely related. However, the phylogenetic clustering methods used in these nomenclature systems are limited by the arbitrariness of setting intra- and inter-cluster diversity thresholds. The lack of a consensus ground truth to define well-delineated, meaningful phylogenetic subpopulations amplifies the difficulties in identifying an informative distance threshold. Consequently, phylogenetic clustering often becomes an exploratory, *ad-hoc* exercise.

Phylogenetic Clustering by Linear Integer Programming (PhyCLIP) was developed to provide a statistically-principled phylogenetic clustering framework that negates the need for an arbitrarily-defined distance threshold. Using the pairwise patristic distance distributions of an input phylogeny, PhyCLIP parameterises the intra- and inter-cluster divergence limits as statistical bounds in an integer linear programming model which is subsequently optimised to cluster as many sequences as possible. When applied to the haemagglutinin phylogeny of HPAI H5Nx viruses, PhyCLIP was not only able to recapitulate the current WHO/OIE/FAO H5 nomenclature system but also further delineated informative higher resolution clusters that capture geographically-distinct subpopulations of viruses. PhyCLIP is pathogen-agnostic and can be generalised to a wide variety of research questions concerning the identification of biologically informative clusters in pathogen phylogenies. PhyCLIP is freely available at http://github.com/alvinxhan/PhyCLIP.

## Introduction

Advances in high-throughput sequencing technology and computational approaches in molecular epidemiology have seen sequence data increasingly integrated into clinical care, surveillance systems and epidemiological studies (Gardy and Loman 2017). Based on the growing number of available pathogen sequences genomic epidemiology has yielded a wealth of information on epidemiological and evolutionary questions ranging from transmission dynamics to genotype-phenotype correlations. Central to all of these questions is the need for robust and consistent nomenclature systems to describe and partition the genetic diversity of pathogens to meaningfully relate to epidemiological, evolutionary or ecological processes. Increasingly, nomenclature systems for pathogens below the species level are based on sequence information, supplementing or even displacing conventional biological properties such as serology or host range (Simmonds et al. 2010; McIntyre et al. 2013). However, existing sequence-based nomenclature frameworks for defining lineages, clades or clusters in pathogen phylogenies are mostly based on arbitrary and inconsistent criteria.

Standardising the definition of a phylogenetic cluster or lineage across pathogens is complicated by differences in characteristics such as genome organization and maintenance ecology. Cluster definitions vary widely even between studies of the same pathogen, limiting generalisation and interpretation between studies as designated clusters, clades and/or lineages carry inconsistent information in the larger evolutionary context (Grabowski et al. 1904; Dennis et al. 2014; Hassan et al. 2017).

In virology, nomenclature systems are largely reliant on absolute distance thresholds that define the maximum genetic divergence tolerated between viruses designated as closely related (Burk et al. 2011; Van Doorslaer et al. 2011; Lauber and Gorbalenya 2012; Donald et al. 2013; Kroneman et al. 2013; Poon et al. 2015; Smith et al. 2015; Poon et al. 2016; Valastro et al. 2016). Groups of closely related viruses are inferred to be phylogenetic clusters when the genetic distance between them is lower than the limit set on within-cluster divergence. Non-parametric distance-based clustering approaches have defined the distance between sequences using pairwise genetic distances calculated directly from sequence data (WHO/OIE/FAO H5N1 Evolution Working Group 2008; Aldous et al. 2012; Ragonnet-Cronin et al. 2013) or pairwise patristic distances calculated from inferred phylogenetic trees (Hué et al. 2004; Prosperi et al. 2011; Poon et al. 2015; Pu et al. 2015; Ortiz and Neuzil 2017). Within-cluster limits on tolerated divergence have been set using mean (WHO/OIE/FAO H5N1 Evolution Working Group 2008), median (Prosperi et al. 2011) or maximum within-cluster pairwise genetic or patristic distance (Ragonnet-Cronin et al. 2013). Some methods incorporate additional criteria, such as the statistical support for subtrees under consideration or minimum/maximum cluster size (Hué et al. 2004; Prosperi et al. 2010; Prosperi et al. 2011; Ragonnet-Cronin et al. 2013). These genetic distance-based clustering approaches are convenient, as they are rule-based and scalable, allowing for relatively easy nomenclature updates. Arguably, flexibility in the distance thresholds allows researchers to curate clusters based on consistency of the geographic or temporal metadata.

The central limitation of approaches based on pairwise genetic or patristic distance is that thresholds to define meaningful within- and between-cluster diversity are arbitrary. For most pathogens, there is no clear definition of a well-delineated phylogenetic unit to underlie nomenclature designation or suggest what additional information would be informative to delineate subpopulations e.g. information on antigenicity or geography or host range. Resultantly, there is no ground truth to optimise distance thresholds when developing a nomenclature system for most pathogens. Partitioning phylogenetic trees into meaningful subsets is therefore complex and is mostly performed *ad hoc* through exploratory analyses with uninformative sensitivity analyses across thresholds. As expected, cluster membership is highly sensitive to the threshold applied and therefore results can be unstable across different cluster definitions (Rose et al. 2017).

There is a need for a consistent, automated and robust statistical framework for determining cluster-defining criteria in nomenclature frameworks. Here, we describe a statistically-principled phylogenetic clustering approach called PhyCLIP. PhyCLIP is based on integer linear programming (ILP) optimisation, with the objective to assign statistically-principled cluster membership to as many sequences as possible. We apply PhyCLIP to the haemagglutinin (HA) phylogeny of the highly pathogenic avian influenza (HPAI) A/goose/Guangdong/1/1996 (Gs/GD)-like lineage of the H5Nx subtype viruses, which underlies the most prominent nomenclature system for avian influenza viruses and which itself is based on a genetic distance approach (WHO/OIE/FAO H5N1 Evolution Working Group 2008).

PhyCLIP is freely available on github (http://github.com/alvinxhan/PhyCLIP) and documentation can be found on the associated wiki page (http://github.com/alvinxhan/PhyCLIP/wiki).

## New approach

PhyCLIP requires an input phylogeny and three user-provided parameters:

i. Minimum number of sequences (*S*) that should be considered a cluster.
ii. Multiple of deviations (*γ*) from the grand median of the mean pairwise sequence patristic distance that defines the within-cluster divergence limit (*WCL*).
iii. False discovery rate (*FDR*) to infer that the diversity observed for every combinatorial pair of output clusters is significantly distinct from one another.

Figure 1A shows the workflow of PhyCLIP which is further elaborated here. First, PhyCLIP considers the input phylogenetic tree as an ensemble of *N* monophyletic subtrees (including the root) that could potentially be clustered as a single phylogenetic cluster, each defined by an internal node *i* subtending a set of sequences *L_i_* (Figure 1B, see Methods). Consequently, as the topological structure of the phylogenetic tree is incorporated in the cluster structure, it is possible to infer the evolutionary trajectory of the output clusters of PhyCLIP if the tree is appropriately rooted. For clarity, we use the term *subtree* to refer to the set of sequences subtended under the same node that could potentially be clustered and the term *cluster* to refer to sequences that are clustered by PhyCLIP within the same subtree.

**Figure 1:**
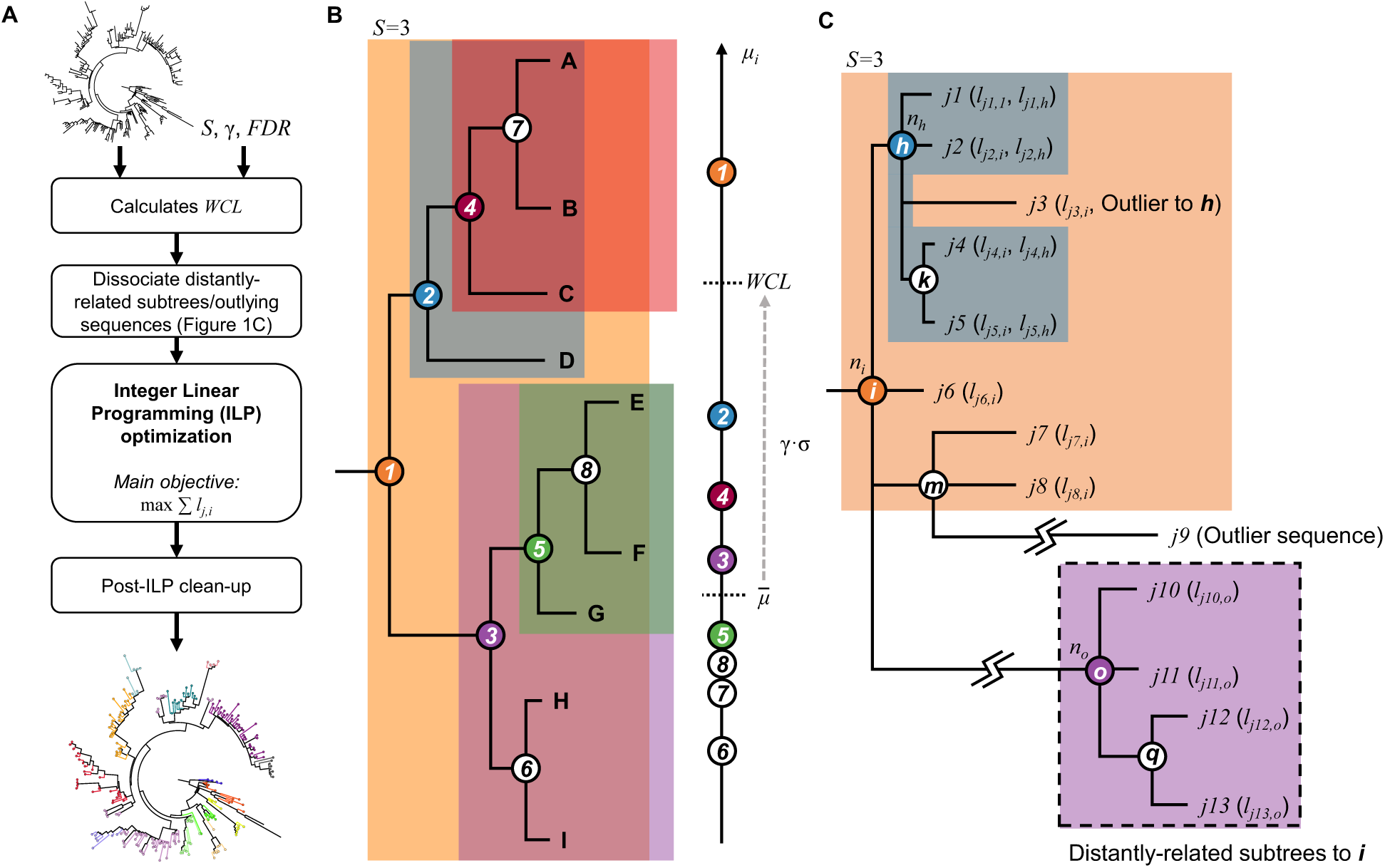
Schematics of PhyCLIP workflow and inference. **A.** Workflow of PhyCLIP. Apart from an appropriately *rooted* phylogenetic tree, users only need to provide *S*, *γ* and *FDR* as the inputs for PhyCLIP. After determining the within-cluster divergence limit (*WCL*), PhyCLIP dissociates distantly related subtrees and outlying sequences that inflate the mean patristic distance (*μ*_i_) of ancestral subtrees. The integer linear programming (ILP) model is then implemented and optimised to assign cluster membership to as many sequences as possible. If a prior of cluster membership is given, this is followed by a secondary optimisation to retain as much of the prior membership as is statistically supportable within the limits of PhyCLIP. Post-ILP optimisation clean-up steps are taken before yielding finalised clustering output. **B.** PhyCLIP considers the phylogeny as an ensemble of monophyletic subtrees, each defined by an internal node (circled numbers) subtended by a set of sequences (letters encapsulated within shaded region of the same colour as the circled number). In this example, only subtrees with ≥ 3 sequences (*S* = 3) are considered for clustering by the ILP model but *WCL* is determined from *μ*_i_ of all subtrees, including the unshaded subtrees 6-8. Only subtrees where *μ*_i_ ≤ *WCL* are eligible for clustering. **C.** Subtrees ***o*** and ***q***, as well as sequence *j9* are dissociated from subtree ***i*** as they are exceedingly distant from ***i***. If sequences *j*1, *j*2, *j*4 and *j*5 are clustered under subtree ***h*** while *j*3 is clustered under subtree ***i*** by ILP optimisation, a post-ILP clean up step will remove *j*3 from cluster ***i***.

The within-cluster internal diversity of subtree *i* is measured by its mean pairwise sequence patristic distance (*μ*_*i*_). PhyCLIP calculates the within-cluster divergence limit (*WCL*), an upper bound to the internal diversity of a cluster, as:

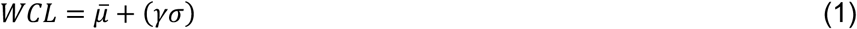

where 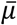 is the grand median of the mean pairwise patristic distance distribution {*μ*_1_, *μ*_2_, …, *μ*_*i*_, …, *μ*_N_} and σ is any robust estimator of scale (e.g. median absolute deviation (*MAD*) or *Qn*, see Methods) that quantifies the statistical dispersion of the mean pairwise patristic distance distribution for the ensemble of *N* subtrees. In other words, only subtrees with *μ*_i_ ≤ *WCL* will be considered for clustering by PhyCLIP (Figure 1B).

### Distal dissociation

The assumption that a cluster must be monophyletic can lead to incorrect assignment of cluster membership to undersampled, distantly related outlying sequences that have diverged considerably from the rest of the cluster (e.g. sequence *j*9 in Figure 1C). These exceedingly distant outlying sequences can also inflate *μ*_i_ of the subtree they subtend, leading to inaccurate overestimation of the internal divergence of the putative subtree. Similarly, distantly related descendant subtrees can artificially inflate *μ*_i_ of their ancestral trunk nodes (e.g. nodes *o* and *q* in Figure 1C). In turn, historical sequences immediately descending from a trunk node *i* will not be clustered if its *μ*_i_ exceeds *WCL* (Figure 1C).

PhyCLIP dissociates any distal subtrees and/or outlying sequences from their ancestral lineage prior to implementing the integer linear programming (ILP) model. For any subtree *i* with *μ*_i_ > *WCL*, starting from the most distant sequence to *i*, PhyCLIP applies a leave-one-out strategy dissociating sequences, and the whole descendant subtree if every sequence subtended by it was dissociated, until the recalculated *μ*_i_ without the distantly related sequences falls below *WCL*. For each subtree, PhyCLIP also tests and dissociates any outlying sequences present. An outlying sequence is defined as any sequence whose patristic distance to the node in question is > 3 × the estimator of scale away from the median sequence patristic distance to the node. *μ*_i_ is recalculated for any node with changes to its sequence membership *L_i_* after dissociating these distantly related sequences. These distal dissociation steps effectively offer PhyCLIP greater flexibility in its clustering construct allowing the identification of paraphyletic clusters on top of monophyletic ones that may better reflect the phylogenetic relationships of these sequences.

### Integer linear programming optimisation

The full formulation of the ILP model is detailed in Methods. Here, we broadly describe how the optimisation algorithm proceeds to delineate the input phylogeny. The primary objective of PhyCLIP is to cluster as many sequences in the phylogeny as possible subject to the following constraints:

i. All output clusters must contain ≥*S* number of sequences.
ii. All output clusters must satisfy *μ*_i_ ≤ *WCL*.
iii. The pairwise sequence patristic distance distribution of every combinatorial pair of output clusters must be significantly distinct from the resultant cluster if sequences from the pair of clusters were to combine. This is the inter-cluster divergence constraint and herein, statistical significance is inferred if the multiple-testing corrected *p*-value for the cluster pair is <*FDR* (see Methods).
iv. If a descendant subtree satisfies (i)-(iii) for clustering (e.g. subtree 5 in Figure 1B) and so does its ancestor, which also subtends the sequences descending from the descendant, (e.g. subtree 3 in Figure 1B), the leaves subtended by the descendant will be clustered under the descendant node (e.g. sequences E, F and G will be clustered under cluster 5 in Figure 1B) while the direct progeny of the ancestor subtree will cluster amongst themselves (e.g. sequences H and I will be clustered under cluster 3 in Figure 1B).

The ILP model is implemented in a third-party linear programming solver fully integrated within PhyCLIP, to obtain the global optimal solution. At the time of this publication, PhyCLIP supports two third-party solvers:

1. Gurobi (http://www.gurobi.com/) is one of the fastest available commercial mathematical programming solvers. Full-featured academic licenses of Gurobi are available for free to users based at any academic institution.
2. GNU Linear Programming Kit (GLPK, http://www.gnu.org/software/glpk) is a popular, free and open-source linear programming solver.

Based on a recent independent benchmark (http://plato.asu.edu/talks/informs2018.pdf), Gurobi outperformed GLPK in both performance and speed (Gurobi solved all 40 Simplex LP test problems while GLPK could only solve 31 of them with a geometric mean runtime that was 52 times longer than Gurobi). As such, it is highly recommended that any users with internet access via an academic domain run PhyCLIP with the Gurobi solver. All clustering results presented in this manuscript were obtained using Gurobi.

### Post-ILP clean-up

While distal dissociation prior to ILP optimisation works well for dissociating distantly related subtrees and sequences, it is ineffective in identifying spurious singletons such as sequence *j*3 in Figure 1C. Here, in terms of sequence patristic distance, sequence *j*3 is an outlying sequence to the descendant node ℎ but not so to the ancestral node *i*. If taxa subtended by subtree ℎ (i.e. *j*1, *j*2, *j*4 and *j*5) were to be clustered without *j*3 which itself is clustered under cluster *i*, PhyCLIP performs a post-ILP optimisation clean-up step that removes *j*3 from output cluster *i*. This is because *j*3 is clearly a topologically outlying taxon to *i* and if unremoved, would imply that sequences clustered under cluster ℎ (i.e. *j*1, *j*2, *j*4 and *j*5) can belong to cluster *i* as well.

PhyCLIP also offers the user an optional clean-up step that subsumes subclusters into their parent clusters if sequences in the descendant subcluster are still associated with the parent cluster (i.e. not removed by distal dissociation) and that coalescing with the parent clusters does not lead to violation of the statistical bounds that define the clustering result. This may be useful if the user prefers a relatively more coarse-grained clustering (e.g. nomenclature building). As mentioned earlier, so long as a statistically significant distinction could be made between a descendant subtree and its ancestral lineage, the ILP model enforces the progeny sequences of the descendant subtree to cluster in the descendant cluster. In turn, PhyCLIP is sensitive to the detection of clusters of highly related or identical sequences that minimally satisfies the minimum cluster size (*S*), as their distributions are statistically distinct from the rest of the population. This sensitivity may lead to over-delineation of the tree and/or multiple nested clusters. Notably, these sensitivity-induced subclusters are not false-positive clusters and meet the same statistical criteria as all other clusters. However, some users may want to subsume these subclusters into parent clusters to facilitate higher level interpretation.

### Optimisation criteria

PhyCLIP’s user-defined parameters can be calibrated across a range of input values, optimising the global statistical properties of the clustering results to select an optimal parameter set. The optimisation criteria are prioritised by the research question, as the clustering resolution and cluster definition are dependent on the question, and therefore the degree of information required to capture ecological, epidemiological and/or evolutionary processes of interest. Users may want a high-resolution clustering result, with the phylogenetic tree delineated into a large number of small, high confidence clusters with very low internal divergence, tolerating a higher number of unclustered sequences. Other users may want a more intermediate resolution, with more broadly defined clusters that are still well-separated but encompass the majority of data in the tree (Figure S1A).

PhyCLIP’s optimisation criteria are agnostic to the metadata of the dataset and include: 1) The grand mean of the pairwise patristic distance distribution and its standard deviation. The grand mean is a measure of the within-cluster divergence and can be optimised to select a clustering configuration with the lowest global internal divergence. 2) The mean of the inter-cluster distance to all other clusters and its standard deviation. This can be optimised to select a clustering configuration with well-separated clusters. 3) The percentage of sequences clustered, which can be optimised to minimise the number of unclustered sequences. 4) The total number of clusters and 5) mean or median cluster size, which can be optimised to select a tolerable level of stratification of the tree.

The ranges of input parameters considered are also dependent on the characteristics of the dataset. The minimum cluster size range considered should be a factor of the size of the phylogenetic tree, whereas the multiple of deviation (*γ*) considered should be a factor of the intra- and inter-cluster distance related to the research question.

Metadata can be incorporated to validate PhyCLIP’s optimisation. The spatiotemporal structure of phylogenies can inform clustering results if within-cluster variation in metadata such as collection times or geographic origin is used as a *post-hoc* optimisation criterion. Within-cluster pairwise geographic distance between the origins of sequences can act as an incomplete ground truth to determine whether a clustering result delineates meaningful clusters if there is a reasonable expectation that clusters are defined by spatial factors. The within-cluster deviation in collection dates can also be included as an optimisation criterion if clusters are expected to be temporally structured.

## Results

To evaluate the utility of PhyCLIP we compared its clustering of the global HPAI H5Nx virus data against the WHO/OIE/FAO nomenclature (WHO/OIE/FAO HN Evolution Working Gr 2009; Smith et al. 2015). The WHO/OIE/FAO H5 nomenclature has been updated progressively since its development in 2007 as new sequences are added to the global phylogeny including updates in 2009 and 2015. The primary analysis of PhyCLIP’s performance was assessed with the full dataset of H5N1 haemagglutinin (HA) sequences included in the WHO/OIE/FAO H5 nomenclature update of 2015 (n=4357), with comparison to the WHO/OIE/FAO clade designation. PhyCLIP was run with different combinations of the parameters varied over the following ranges: a minimum cluster size of 2-10, a multiple of deviation (*γ*) of 1-3, and an FDR of 0.05, 0.1, 0.15 or 0.2. The optimisation criteria were prioritised as follows: 1) percentage of sequences clustered, 2) grand mean of within-cluster patristic distance distribution, 3) mean within-cluster geographic distance and 4) mean of the inter-cluster distances.

The percentage of sequences clustered was prioritised as the primary optimisation criterion to ensure that the maximum number of sequences were assigned a nomenclature identifier. Mean within-cluster geographic distance was included as a *post-hoc* optimisation criterion as many avian influenza viruses cluster with high spatial consistency owing to their transmission dynamics in localised avian populations. For influenza viruses endemic to poultry such as H5Nx, this is likely owing to increased local transmission during outbreaks in large poultry populations, as well as the associated sampling biases (Smith et al. 2015). Within-cluster genetic divergence was optimised with higher priority than within-cluster mean geographic distance, as the use of phylogenetic geographic structure as a ground truth for avian influenza viruses is restricted by the long-distance dissemination of related viruses through mechanisms such as the poultry trade or migration of wild birds (WHO/OIE/FAO H5N1 Evolution Working Group 2014a; Smith et al. 2015a). The within-cluster geographic distance was calculated for each cluster in each clustering result as the mean within-cluster pairwise Vicenty distance in miles.

The temporal consistency of clusters can also be used as optimisation criteria for measurably evolving viruses such as Influenza A virus (Drummond et al. 2003). Results ranking the grand mean within-cluster standard deviation in sampling dates of each clustering result as the fourth optimisation criterium, with mean of the inter-cluster distance in fifth, were identical to those only including the above mentioned four optimisation criteria.

As PhyCLIP incorporates topological information of the phylogeny into the clustering construct, non-terminal internal nodes with zero branch lengths can lead to erroneous clustering and over-delineation (Figure S1B). Such internal nodes are usually found in bifurcating trees as representations of polytomies, arising from a lack of phylogenetic signal among the sequences subtended by the node to resolve them into dichotomies. As such, prior to implementing PhyCLIP, all non-terminal, zero branch length nodes in the input phylogenetic trees were collapsed into polytomies, which more accurately depicts the relationship between identical/indiscernible sequences and/or ancestral states. In the H5Nx analysis, all subclusters were subsumed if the statistical requisites of the parent clade were maintained, to aid in easing the interpretation of the nomenclature designation (as discussed in the New Approach section).

### Influence of the parameters

The influence of the parameters on PhyCLIP’s clustering properties was assessed with the 2015-update H5 phylogeny. Lower multiples of deviation (*γ*) define a more conservative expected range for tolerated within-cluster divergence, informed by the global pairwise patristic distance distribution (Figure S2). As a result, clusters designated at a *γ* of 1 have the lowest internal divergence, measured by the grand mean of the pairwise patristic distance distribution (Figure 2C). These clusters are expected to be highly related, with low variation in clustered sequence spatiotemporal metadata (Figure 2E-F). More conservative ranges of tolerated within-cluster divergence result in a higher clustering resolution with a greater number of clusters, lower mean cluster sizes and a higher percentage of sequences unclustered (Figure 2A-B). A higher *γ* increases the limit of tolerated within-cluster divergence, resulting in a lower clustering resolution that coalesces smaller clusters into larger, more internally-divergent clusters. The collapsing of the smaller clusters decreases the total number of clusters while concurrently increasing the percentage of sequences clustered and mean cluster size. The influence of *γ* is less pronounced for the mean inter-cluster distance, with no apparent distinction between *γ* = 1 and 2. The total number of clusters decreases approximately linearly as the minimum cluster size (*S*) increases from two towards ten (Figure 2A). Lower FDRs are more conservative in designating the pairwise patristic distance distributions of two clusters as statistically distinct. A higher or less conservative FDR therefore designates more similar distributions as distinct from one another, increasing the number of clusters (Figure 2A). The effect of FDR is muted at a higher minimum cluster size or higher *γ*, as these parameters designate larger clusters, which limits the number of clustering configurations available.

**Figure 2:**
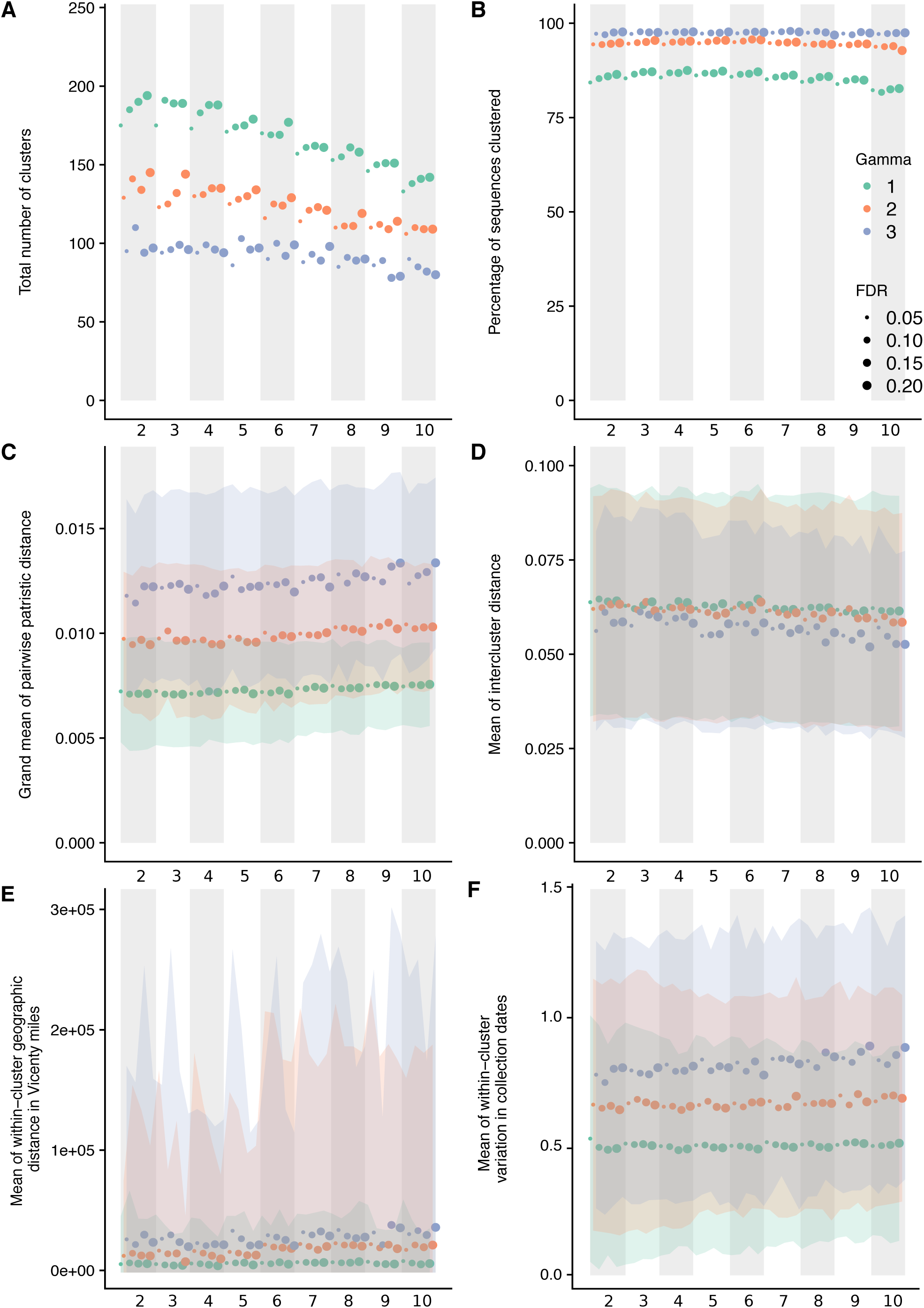
Influence of parameters on the clustering properties of PhyCLIP in the WHO/OIE/FAO 2015-update phylogeny. Figures A-F have the parameter set combinations ordered according to minimum cluster size, FDR and *γ* on the x axis. The banded background and x-axis subscript numbering indicate the minimum cluster size of the parameter set. Marker colour and size is indicative of the *γ* and the FDR respectively of the parameter set as indicated by the legend in Figure B. **A**. Total number of clusters. **B**. Percentage of sequences clustered. **C**. Grand mean of the pairwise patristic distance distribution. **D**. Mean of the inter-cluster distance to all other clusters. **E**. Mean within-cluster geographic distance calculated in Vicenty miles. **F**. Mean within-cluster standard deviation in collection dates.

### Optimal PhyCLIP clustering result for HPAI avian H5 viruses

For the full phylogeny of Gs/GD-like H5 viruses from the 2015 nomenclature update, the optimal parameter set combined a minimum cluster size of 7, an FDR of 0.15 and a *γ* of 3. The optimal clustering configuration clustered 98% of the sequences into a total of 89 clusters with a median cluster size of 21 sequences. The topology of the optimal clustering result yields informative source-sink trajectories that are supported by previously reported phylogenetic and phylogeographic evidence of the global panzootic of the Gs/GD-like H5N1 lineage (Duan et al. 2008; Wang et al. 2008; Smith et al. 2015; The Global Consortium for H5N8 and Related Influenza Viruses 2016).

Principally, pathogen nomenclature systems should delineate population structure, highlighting the underlying population dynamics that may be informative about the evolutionary trajectory of pathogen variants. The distal dissociation approach of PhyCLIP produces a clustering topology where divergent subclusters nest within a larger cluster structure termed a supercluster, as exemplified with WHO/OIE/FAO clade 2.1x viruses in Figure 3. Sufficiently diverse subclusters are dissociated from the ancestral trunk node of a putative cluster. This enables the remaining sequences that meet the statistical criteria to cluster with the ancestral node based on their pairwise patristic distance, as the divergent subcluster is no longer inflating the ancestral node’s mean pairwise patristic distance above the within-cluster limit. Cluster A in Figure 3 depicts the supercluster topology: the source population viruses (tips in yellow) are annotated as A, and the divergent descendant subclusters are annotated as A.1, A.2 and A.3 respectively. This approach captures source-sink ecological dynamics: the supercluster acts as a putative source population to its subclusters, reflecting the clear evolutionary divergence and trajectory of descendants of the source population (sub-lineages). The nomenclature system algorithmically imposed on PhyCLIP’s clustering for avian influenza is designed to enhance the evolutionary information in the clustering (see Methods).

**Figure 3:**
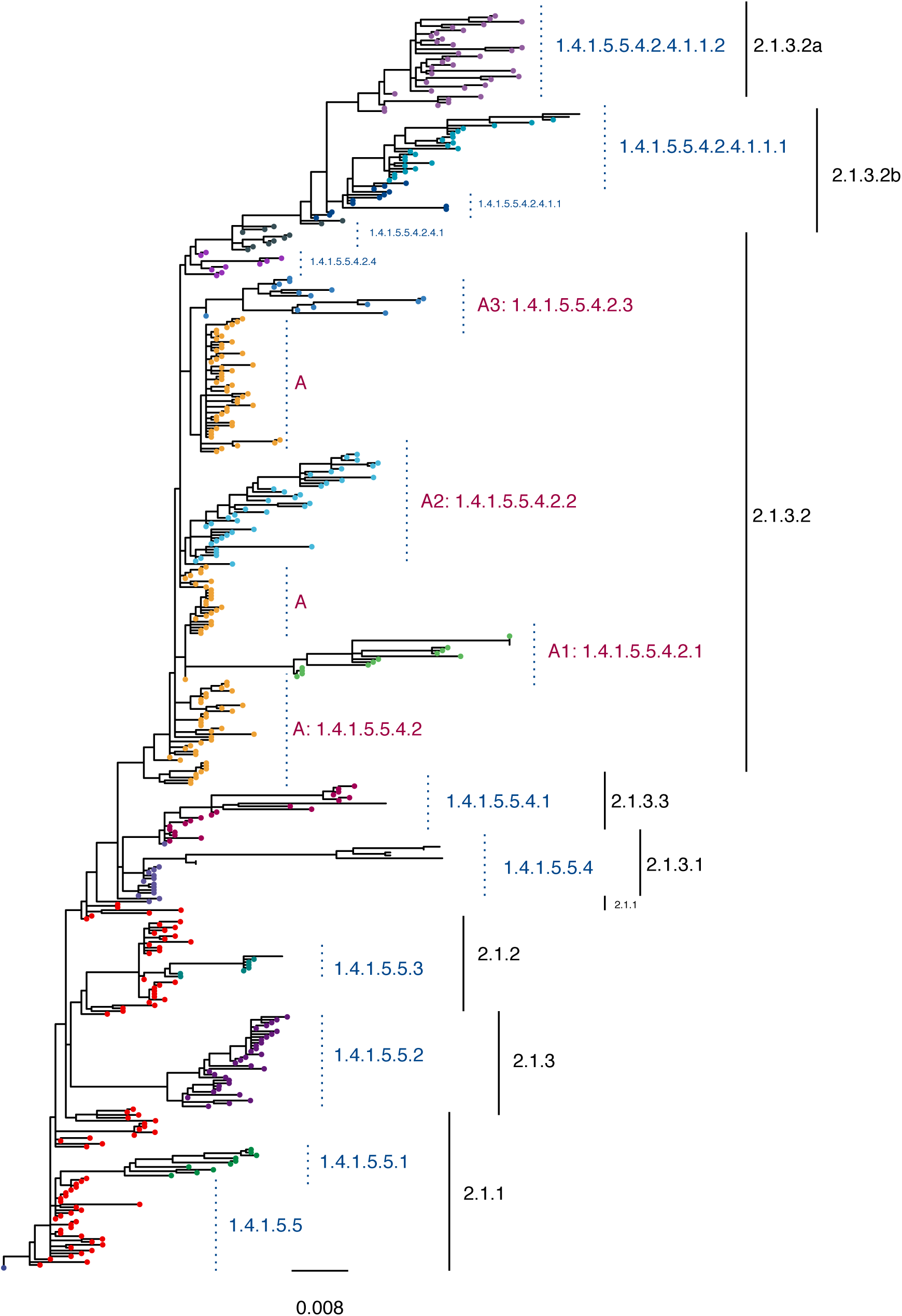
Phylogeny of the Clade 2.1x viruses circulating in Indonesia. The WHO/OIE/FAO H5 nomenclature is annotated in black. PhyCLIP’s cluster designation is indicated in blue, corresponding to tip colour. PhyCLIP’s supercluster topology is exemplified by Cluster A. The source population of the supercluster is annotated as A in pink, with tips coloured yellow. The divergent descendant clusters are annotated as A.1, A.2 and A.3 respectively here. The letter A here is shorthand for its nomenclature address, 1.4.1.5.5.4.2. This nomenclature address indicates that supercluster A is the second descendant of cluster 1.4.1.5.5.4 (indicated in light purple), which in turn is the forth descendant of the source supercluster 1.4.1.5.5, indicated in red. See Methods sections for full explanation of nomenclature addresses.

PhyCLIP’s optimal cluster designation delineated the spatiotemporal structure of the phylogeny at high resolution (Figure S3). Viruses circulating in south, central and northeast China and Hong Kong in 1996-2003 acted as the source population for the emergence of the classical viruses, seeding four lineages (cluster 1, seeding cluster 1.1-1.4, Table S1). The second supercluster captures the first major wave of expansion into neighbouring countries in east and southeast Asia in the early 2000s, with a source population of viruses circulating in south central, east and north China, Viet Nam and Hong Kong in 2000-2003 (1.4 and 1.4.1 and their descendant lineages). The third supercluster captures the second major wave of expansion of the Gs/GD-like H5 viruses, characterised by global spread (cluster 1.4.1.5 and its descendants). The source population of viruses from east, south central and southwest China, Hong Kong and Viet Nam circulated from 2002-2005, giving rise to diverse and distinct viral lineages in different regions globally (1.4.1.5.1-6). The supercluster topology highlights single lineage introductions for countries with endemic circulation such as Indonesia and Egypt, but delineates multiple co-circulating lineages structured over time. The clustering topology also highlights multiple incursions of diverse viruses into countries such as South Korea and Japan (Table S3).

In addition to source-sink dynamics, distal dissociation also identifies probable outlying sequences, defined as sequences more than 3 times the estimator of scale away from the median patristic distance to the node. For example, PhyCLIP identifies seven outliers in its delineation of WHO/OIE/FAO clade 2.3.2.1c in the 2015 phylogeny (indicated by red tip-points in Figure 4). These sequences may represent under-sampled populations with unobserved diversity, introductions from otherwise unsampled populations or lower quality sequences introducing error into phylogenetic reconstruction.

**Figure 4:**
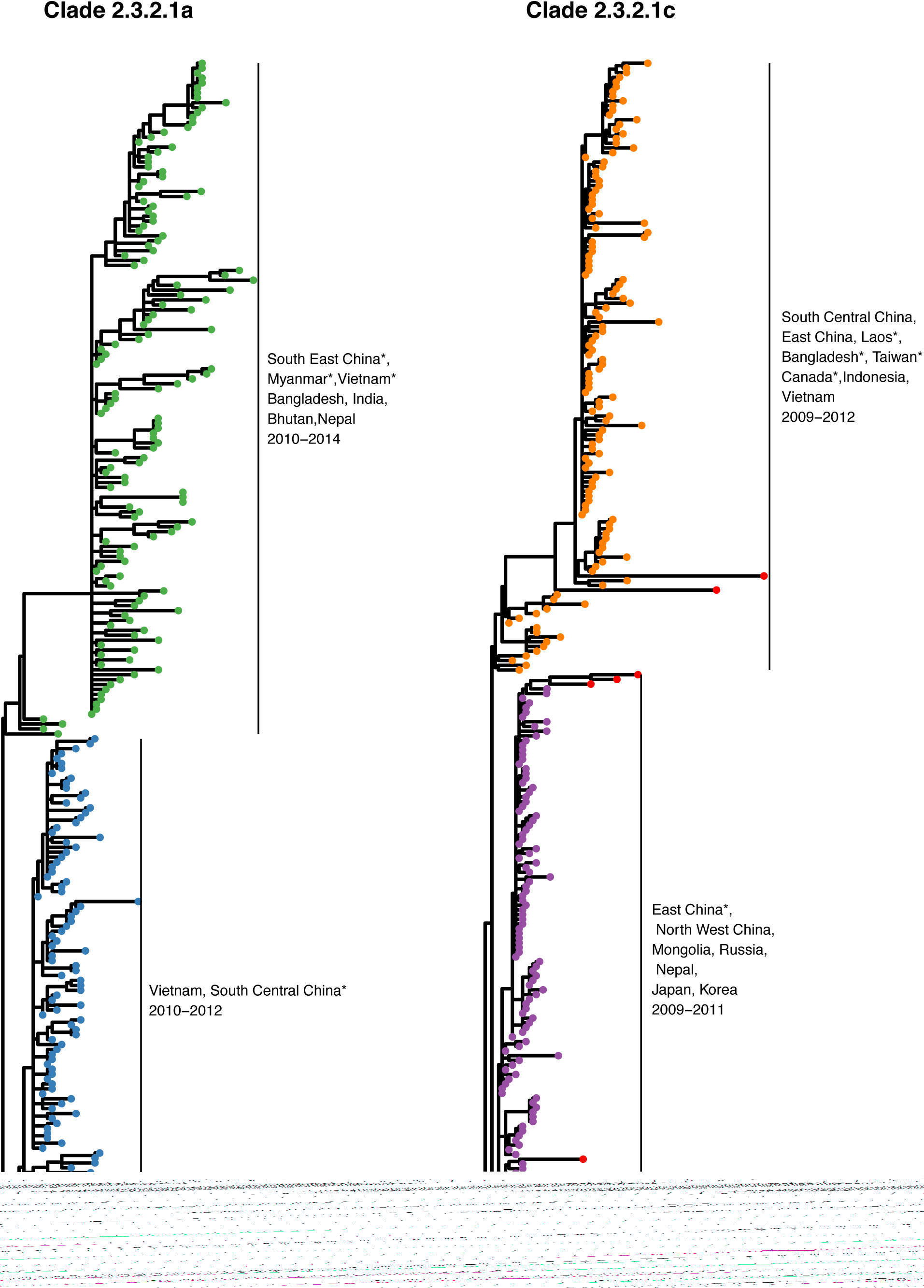
PhyCLIP’s delineation of WHO/OIE/FAO demarcated clades 2.3.2.1a (A) and 2.3.2.1c (B). Tips are coloured according to PhyCLIP’s cluster designation. The tips coloured in red in B are viruses that were designated as outliers by PhyCLIP’s outlier detection. Countries represented by single viruses in the cluster are indicated with an asterisk.

### Comparison to the WHO/OIE/FAO H5 nomenclature

The current WHO/OIE/FAO nomenclature system designates 43 different clades and 7 clade-like groupings for the full H5 phylogeny as of the 2015 update (Smith, Donis, and WHO/OIE/FAO H5 Evolution Working Group 2015) (Table S2). PhyCLIP recovers the current WHO/OIE/FAO H5 nomenclature with varying degrees of agreement across parameter sets, as measured by the variation of information (VI) between the clustering partitions (Figure S4). VI is an information theoretic criterion for comparing partitions of the same data set, based on the information lost and gained when moving between partitions (Meilă 2007). A lower VI indicates more similar partitions. Parameter sets with a *γ* of 3 consistently had the lowest VI compared to the WHO/OIE/FAO system, indicating that the WHO/OIE/FAO nomenclature system has the highest agreement with PhyCLIP clustering results that tolerate higher within-cluster divergence.

In the optimal clustering result, PhyCLIP delineates the spatiotemporal structure of the phylogeny with a higher resolution than the WHO/OIE/FAO nomenclature system (89 vs 50 phylogenetic units, Figure S3). The supercluster structure of the PhyCLIP clustering topology recapitulates the hierarchical structure of the WHO/OIE/FAO nomenclature (Figure 3). Simultaneously, PhyCLIP’s clustering captures clear lineage distinctions for viruses from different geographic regions and years in several WHO/OIE/FAO demarcated clades. For example, PhyCLIP delineates clade 2.3.2.1a into two separate clusters: 1) a cluster that circulated in Viet Nam in 2011-2012, with sporadic detection in south central China and 2) a cluster that circulated largely in Bangladesh, India, Bhutan and Nepal from 2010 to 2014, with single viruses detected in south east China, Viet Nam and Myanmar (Figure 4A). PhyCLIP also delineates clade 2.3.2.1c into two clusters: 1) a cluster that captures the expansion of viruses from north west and east China into Mongolia, Russia, Nepal, Japan and Korea for the period 2009-2011, and 2) a cluster that predominantly circulates in China, Viet Nam and Indonesia for 2009-2012, with single viruses from Lao PDR, Bangladesh and Taiwan (Figure 4B).

### Impact of sampling

PhyCLIP’s clustering results are sensitive to the diversity in the input population that informs the global distribution and resultantly sampling. The influence of sampling was assessed by comparing the optimal clustering result of the phylogeny underlying the WHO/OIE/FAO H5 2015 nomenclature (n=4357) to the phylogeny underlying the 2009 nomenclature update (n=1224), a subset nested in the 2015-update phylogeny. The WHO/OIE/FAO 2009 nomenclature update was performed after the geographic expansion and divergence of clade 2.2, which necessitated further delineation into clade 2.2.1. It designated 20 clades, including 8 third order clades (WHO/OIE/FAO HN Evolution Working Gr 2009). The WHO/OIE/FAO 2015 nomenclature update includes approximate 3.5-times the number of sequences as the 2009 nomenclature update, and includes novel clade designation to the fourth and fifth order WHO/OIE/FAO H5 Evolution Working Group 2015). The optimal PhyCLIP parameter set for the 2009 WHO/OIE/FAO nomenclature system combines a minimum cluster size of 3, a FDR of 0.2 and a *γ* of 3. In the 2009 tree, this clustered 98% of the n=1224 viruses into 39 clusters, with a median cluster size of 12 (Figure S5).

Overall, the source-sink inference of PhyCLIP’s clustering topology is largely consistent between the WHO/OIE/FAO 2009 and 2015 update phylogeny optimal clustering results (Table S1). The optimal result for the 2009 update phylogeny captures a similar topology and source population for the South East Asian (clusters 1.3.1 and 1.3.1.1) and the post-2005 global wave of expansion (cluster 1.3.1.1.2.2.2) compared to the optimal 2015 clustering, with substantial overlap between the source populations identified (100% and 83% for source populations for southeast Asia wave and global wave respectively).

Changes in the clustering topology between the 2009 and 2015 update phylogenies are expected as the underlying datasets are substantially different. More than 3000 viruses were added to the tree in the six years between nomenclature updates. The Gs/GD-like H5 viruses evolved significantly in the intervening period owing to genetic drift and reassortment. The addition of a large number of divergent viruses to the underlying dataset fundamentally alters the ensemble statistical properties of the tree, driving changes in the clustering configuration by changes in the global patristic distance distribution, topology and statistical power between datasets. As a result, the ecological inferences drawn from the 2015 clustering topology are different from that of the 2009 phylogeny (Table S1).

Primarily, the addition of a set of highly divergent sequences increases the spread of the global pairwise patristic distance distribution (Figure S2). The within-cluster limit it informs increases concurrently, increasing the tolerance of allowable within-cluster divergence. In the distal dissociation approach, increased tolerance of divergence would allow for the incorporation of more distant trunk viruses into supercluster source populations if the enclosed viruses are sufficiently distinct to be dissociated as independent clusters (Figure S6). If the within-cluster limit is lowered, inclusion of the considered trunk viruses will violate the within-cluster limit. Resultantly, these trunk viruses and their descendants will be assessed for clustering as independent subtrees.

Clustering changes between 2009 and 2015 update phylogenies are also induced by the local effects of the addition of multiple lineages to the 2015 phylogeny within clusters defined in 2009 owing to their continued circulation and diversification post-2009. Notably, many distinct clusters in the 2009 phylogeny are structured as source populations in superclusters in the 2015 phylogeny (Figure S7). Here, PhyCLIP identifies that the statistical properties of these divergent post-2009 lineages are distinct enough to reliably dissociate them from the ancestral node and delineate them as separate clusters. The viruses present in the 2009 phylogeny that these divergent lineages descend from meet the within-cluster limit after the dissociation and are structured as the source population to the post-2009 nested diversity.

Topological differences between phylogenetic trees built from different underlying datasets can also drive changes in PhyCLIP’s clustering, as observed for the classical clade 0 viruses (Figure S6). The source population of the classical clade viruses for both the 2009 and 2015 updates optimal clustering result is estimated to have originated from south central and east China and Hong Kong in 1997-2003. However, the 2015 cluster designation resolves an additional seed lineage within the 2009-source population (Figure S6). In the 2009 phylogeny, this additional cluster forms part of the source population as it is part of the trunk of the tree. The equivalent cluster does not form part of the trunk of the tree in the 2015 phylogeny and is dissociated as a statistically distinct cluster. Moreover, the substantial increase in the number of viruses between the 2009 and 2015 datasets along with the increase in diversity results in more statistical power to delineate among groups of viruses resulting in a higher clustering resolution for the 2015 phylogeny.

### Comparison of optimal to suboptimal clustering results

So far, we have focused our interpretation on the optimal PhyCLIP clustering. To ensure that our results were robust across similarly optimal PhyCLIP parameter sets we compared the optimal set against the next four similarly optimal sets. Comparing the top 5 clustering results ranked by the optimality criterion (in order of greatest number of sequences clustered, lowest internal genetic and geographic divergence, and greatest average between-cluster distance), the clustering result from the optimal parameters set of the 2015 phylogeny was generally consistent with those generated from the four highest-ranked suboptimal parameter sets (see Figure S8). Each of the top four suboptimal clustering was found to have low VI (0.817-0.984) relative to the optimal clustering, with a large proportion (74.4%-82.7%) of viruses clustered in the same corresponding clusters. The supercluster source populations leading to the early 2000 expansion into east and southeast Asia as well as the global expansion in 2005 were similarly found in all suboptimal results.

However, changes to parameter sets fundamentally changed the statistical constraints defining the clustering solution space and in turn, altered the partitions between resultant clusters. Specifically, in this case where *γ* = 3 in all five optimal/suboptimal parameter sets, varying minimum cluster size not only changed the distribution of putative subtrees for clustering but the distribution of inter-cluster divergence *p*-values for multiple-testing correction as well. As such, while the global superclusters were largely recapitulated in the suboptimal results, local partitions of co-circulating viruses descending from these supercluster sources, and consequently the inferences of source-sink dynamics, varied amongst the different parameter sets.

### PhyCLIP clustering of the 1996-2018 H5Nx phylogeny

In recent years the Gs/GD-lineage of H5 viruses has undergone substantial evolution, with viruses from WHO/OIE/FAO clade 2.3.4.4 reassorting with co-circulating viruses to give rise to multiple H5Nx subtypes including H5N2, H5N5, H5N6 and H5N8. We applied PhyCLIP to a phylogeny representing the Gs/GD-lineage up to and including early 2018 to investigate how the global expansion of the H5Nx viruses changes clustering inference (n=7898) (Figure S9, S10). Applying the same optimisation approach described above, the optimal parameter set for the 2018 phylogeny combines a minimum cluster size of 4, a FDR of 0.2 and a *γ* of 3. This parameter set clustered 97% of the viruses into 135 clusters, with a median cluster size of 23 (Figure S11).

The addition of the H5Nx viruses collected from 2014-2018 to the 2015 phylogeny changed the distribution in two ways: 1. it added diversity to the right tail of the distribution, owing to the increased divergence of the H5Nx viruses compared to the H5N1 viruses; 2. it increased the number of putative clusters with low internal divergence, as a large amount of the H5Nx viruses possess highly similar HA genes owing to both sampling biases during outbreaks and the relative short circulation time following their emergence. This shift in the distribution reduced the within-cluster limit compared to that of the 2015 dataset (Figure S2).

Filtering the 2015-update and 2018 datasets (see Methods) resulted in changes in tree topology and overall sequence diversity, and consequently altered the ecological inference of source-sink clusters circulating from 1997-2005 (Table S1). However, the ecological inferences of the second major wave of expansion, the post-2005 global expansion characterised by cluster 1.2.1.1.1.3.2 and its descendants 1.2.1.1.1.3.2.1-8, were largely consistent across the 2009 (cluster 1.3.1.1.2.2.2), 2015 (cluster 1.4.1.5) and 2018 (cluster 1.2.1.1.1.3.2) trees, including a shared core source population (Table S1).

The WHO/OIE/FAO clade 2.3.4.4 viruses are of interest owing to their reassortment promiscuity and rapid global expansion. PhyCLIP delineates the clade 2.3.4.4 viruses into two distinct lineages, seeded from a source population of viruses circulating in east and south-central China and Malaysia in 2005-2010 (cluster 7.8, Table S1). The first lineage circulated in east, south central and northeast China from 2008 to 2011 (7.8.2, Figure S11, Table S1). The second lineage (7.8.3) circulated in south central and east China in 2008-2012 and seeded six distinct sub-lineages: Lineage 7.8.3.1 circulated in China from 2010 to 2014 before expanding to Viet Nam and circulating there for 2014-2015. Lineage 7.8.3.2 captures the global expansion of viruses from 2009 onwards. This includes the early subclade of H5N8 viruses described in Lycett et al (The Global Consortium for H5N8 and Related Influenza Viruses 2016). Lineage 7.8.3.3 was restricted to China and was detected in 2013-2016. Lineage 7.8.3.4 also captures a pan-national lineage that was detected from 2014 to 2016, and captures the more recent H5N8 subclade described in Lycett et al (The Global Consortium for H5N8 and Related Influenza Viruses 2016). Lineage 7.8.3.5 circulated in east and southeast Asia from 2013 to 2017. Lineage 7.8.3.6 is seeded from a source population of viruses circulating in east and southeast Asia, expanding into multiple co-circulating H5N6 southeast Asian lineages from 2013 onwards (Table S1).

### Benchmarking against other phylogenetic clustering tools

PhyCLIP was benchmarked for performance against two open-source non-parametric clustering tools, PhyloPart (Prosperi et al. 2011) and ClusterPicker (Ragonnet-Cronin et al. 2013). Both tools require a phylogenetic tree as input, as well as a user-specified distance threshold and minimum statistical node-support level. Additionally, both algorithms carry out a depth-first traversal of the tree, considering subtrees as putative clusters if the node support is above the user-defined level. In PhyloPart, the user specifies a percentile of the global pairwise patristic distance distribution as a threshold. If the median of the pairwise patristic distances of the putative cluster is below the percentile threshold, a cluster is designated. ClusterPicker requires a user-defined maximum pairwise genetic distance (calculated as p-distance directly from the sequences) threshold for cluster designation. In both tools, a subtree is designated as a cluster if it meets the respective clustering criteria. If the subtree violates the clustering criteria, the algorithm tests the children of the subtree as potential clusters until a leaf is reached, when no cluster is designated in the path.

In contrast, traversal order has no bearing on the clustering outcomes of PhyCLIP. Although PhyCLIP parses the input phylogeny by level-order, prior to ILP optimisation, PhyCLIP dissociates outlying taxa if *μ*_i_ < *WCL* and proceeds with full distal dissociation heuristics described in the New Approach section if otherwise for every internal node *i* in the input tree. In both cases, tip dissociation is performed by ranking taxa based on their patristic distance to node *i* (i.e. the common ancestor) without consideration of their topological placement. Finally, all putative subtrees (i.e. tree nodes) after distal dissociation are given equal consideration by ILP optimisation to maximally assign cluster membership to all tips (see New Approach). In doing so, not only does PhyCLIP allow for paraphyletic clustering, tree traversal order does not affect clustering results.

Accepted practice for these tools is to incorporate previous knowledge of sequence divergence into a distance threshold or to calibrate the threshold over a tolerable range with metadata or expert consensus. The two methods were applied to the 2009-update phylogeny (n=1224 sequences) with thresholds ranging from 0.005 to 0.05 substitutions/site. For PhyloPart, the respective percentile of the global pairwise patristic distance distribution was chosen to match the distance threshold. Required bootstrap support level was set to 0 in both methods to make it comparable to PhyCLIP, which lacks node-support criteria. The optimal threshold was selected by maximisation of the mean silhouette index across the clustering partitions (see Methods). All programs were run on the Ubuntu 16.04 LTS operating system with an Intel Core i7-4790 3.60 GHz CPU.

The optimal thresholds and clustering statistics for each of the methods are reported in Table S4. A direct comparison of cluster inference between PhyCLIP and the other methods is difficult owing to notable differences in cluster definitions, as these methods were largely designed to detect highly related clusters of sequences linked by direct transmission events. The optimal clustering result for ClusterPicker by silhouette maximisation had a very low maximum genetic distance threshold at 0.5%. (Figure S12). This resulted in a highly stratified tree with 246 small, highly-related clusters and 33.8% outliers, compared to PhyCLIP’s 39 clusters and 2% outliers (VI to PhyCLIP of 2.7) (Figure S13, Table S4).

Clustering results between PhyCLIP and PhyloPart’s optimal results showed better correspondence, with PhyloPart designating 37 clusters to PhyCLIP’s 39 (VI to PhyCLIP of 0.64, Figure S13, Table S4). However, the cluster delineations and inferences drawn are substantially different between the two methods (Table S5). The nomenclature scheme developed for PhyCLIP was applied to PhyloPart’s optimal clustering result for a more meaningful comparison. PhyCLIP’s distal dissociation approach allows for the identification of paraphyletic clusters, forming supercluster topologies throughout the tree (as discussed above). Notably, PhyloPart’s depth-first approach and monophyletic cluster criteria prevent it from designating paraphyletic clusters, obscuring the suggestive source-sink dynamics of H5N1’s expansion wave identified by PhyCLIP’s distal dissociation approach (Table S5). Concurrently, PhyloPart is unable to identify hierarchical clusters, which PhyCLIP identifies as divergent trajectories nested in larger clusters (Figure S13).

PhyCLIP is appreciably more computationally intensive than PhyloPart and ClusterPicker as it not only parses the global pairwise patristic distance distribution of the phylogeny but recursively recalculates the distribution for subtrees in the distal dissociation approach, performs hypothesis testing across every combinatorial pair of subtrees to test their inter-cluster divergence, as well as optimise the ILP model. To relieve some of the computational cost, PhyCLIP is written in Python 2.7 employing multiprocessing modules to parallelise the computational tasks involved resulting in ~3.2x times speedup with 8 CPU cores relative to a single core run (Table 1).

**Table 1.** Benchmarking the performance of PhyCLIP against widely-used phylogenetic clustering tools

| Approach | Time to completion | Peak memory usage | Number of CPUs |
| --- | --- | --- | --- |
| PhyCLIP | 1 hour 4 minutes | 2.0 GB | 8 |
|  | 3 hours 25 minutes | 1.7 GB | 1 |
| ClusterPicker | 2.8 minutes | 0.3 GB | 1 |
| PhyloPart | 10.6 minutes | 4.1 GB | 8 |

Despite the differences in computation time, the principal advantage of PhyCLIP is its use of the background genetic diversity to inform its within-cluster limit without the need to arbitrarily define it or calibrate it over a range of thresholds. This is especially helpful as there is typically a lack of prior knowledge on meaningful delineation of phylogenetic units for most pathogens to recommend a range of distance thresholds. Additionally, PhyCLIP’s distal dissociation and outlier detection approaches are capable of identifying informative paraphyletic and hierarchical clusters, unlike the other tools.

## Discussion

PhyCLIP provides a statistically-principled, phylogeny-informed framework to assign cluster membership to taxa in phylogenetic trees without the introduction of arbitrary distance thresholds for cluster designation. PhyCLIP uses the pairwise patristic distance distribution of the entire tree to inform its limit on within-cluster internal divergence against the background genetic diversity of the population included in the phylogeny. Testing against the global background genetic diversity indicates whether the putative clustered sequences are sufficiently more related to one another than to the rest of the dataset to be designated a distinct cluster.

PhyCLIP’s cluster assignment is agnostic to metadata but is capable of capturing the geographic and temporal structure of the H5 phylogeny informatively. PhyCLIP recovers the overall structure of the current WHO/OIE/FAOH5 nomenclature developed on a sequence divergence threshold but delineates more informative, higher resolution clusters that capture geographically-distinct subpopulations. PhyCLIP therefore plausibly provides the foundation for an alternative nomenclature that minimises the limitations of currently employed approaches.

PhyCLIP’s clustering is expected to improve with the addition of new sequences to the tree as new information about the genetic diversity and evolutionary trajectory of the pathogen becomes known and can be incorporated into the background diversity of the tree that informs the algorithm. Additionally, topological information that captures how sequences are related by common ancestors is inherently incorporated in PhyCLIP owing to its distal dissociation approach. The distal dissociation approach also does not assume all clusters are monophyletic as the most recent common ancestor of all tips in a cluster is not assumed to have any descendants. As such, PhyCLIP can identify nested clusters both as clusters with sufficiently high information content to meet the statistical requirements of cluster designation or sufficiently diverse clusters that are dissociated from their ancestral nodes. The designation of divergent descendant clusters nested within a supercluster suggestively captures source-sink population dynamics that may be informative about the evolutionary trajectory of the clustered sequences. At the same time, users could also opt for PhyCLIP to subsume subclusters that do not violate the statistical criteria of the parent clusters into the latter, aiding higher level interpretation. Importantly, the distal dissociation approach also identifies highly divergent outlying sequences that may be indicative of under-sampled diversity.

For pathogens that evolve more rapidly than they spread geographically, it is expected that clusters of related sequences would be temporally structured. However, it is important to consider the distribution of sampling times, which can drive clustering artificially. This is especially pertinent for transmission dynamic studies, where clustering is often driven by heterogeneity in sampling rates across subpopulations rather than heterogeneity in transmission rates (Poon 2016; McCloskey and Poon 2017). PhyCLIP can be applied to time-resolved phylogenies in heterochronous datasets. However, molecular clock analyses make strong biological assumptions and require sufficient temporal signal to inform the model reconstructing the statistical relationship between genetic divergence and time (Rambaut et al. 2016). These models rely on high-quality sampling dates and alignments free of sequence error and laboratory-altered strains or recombinant viruses to reconstruct valid and unbiased time-scaled phylogenies (Rambaut et al. 2016). As PhyCLIP centrally operates on the branch lengths of the phylogeny, we recommend it is only applied to robust time-resolved phylogenies after a thorough investigation of the temporal signal as well as a rigorous assessment of model and prior assumptions (Boskova et al. 2018).

PhyCLIP’s methodology has limitations. Notably, PhyCLIP is tree-based and is therefore subject to error in phylogenetic reconstruction. PhyCLIP does not include criteria for the statistical support of nodes under consideration, which omits uncertainty in phylogenetic reconstruction. However, high statistical support for a node does not necessarily indicate that all sequences subtended by it are highly related but merely reflects the statistical support of the bipartition to the exclusion of other sequences. Additionally, the relationship between the statistical significance of internal nodes and population dynamics is unresolved as is an appropriate definition of a robustly supported node (Zharkikh and Li 1992; Susko 2009; Anisimova et al. 2011; Kumar et al. 2012; Volz et al. 2012). There is often less phylogenetic signal to resolve internal nodes subtending small subtrees in measurably evolving populations, increasing uncertainty in the arrangement of the internal structure of smaller subtrees. If a statistical support threshold is set for nodes, these viruses will consistently be left unclustered or will be forced to coalesce with more ancestral nodes subtending larger clusters, which would violate PhyCLIP’s statistical framework.

As with any phylogenetic clustering methods, PhyCLIP is also sensitive to variation in sampling rates (Volz et al. 2012). There is a significant surveillance bias towards certain pathogens (e.g. HPAI H5) owing to their consequences for animal and human health. The evolution and divergence of these pathogens are currently captured in surveillance data as a more accurate approximation to a continuum of evolution. PhyCLIP’s clustering is strongly influenced by the diversity in the input population it tests against and will perform best when the background diversity of the phylogeny is complete or representative.

Clusters identified by PhyCLIP should not be interpreted as sequences linked by rapid direct transmission events. Transmission dynamic studies aim to integrate epidemiological clustering with phylogenetic clusters to study transmission chains or local outbreak networks by assuming putative transmission links between highly related sequences (Hassan et al. 2017). Datasets from transmission dynamic studies are likely to be sampled from localised outbreaks over a very specific period of time. The global distribution generated from the resulting phylogenetic trees will not contain sufficient information or power to meaningfully compare subpopulations to identify high confidence transmission clusters.

In conclusion, PhyCLIP provides an automated, statistically-principled framework for phylogenetic clustering that can be generalised to research questions concerning the identification of biologically informative clusters in pathogen phylogenies.

## Materials and methods

### Robust estimator of scale (deviation)

PhyCLIP computes the robust estimator of scale (σ) either as the median absolute deviation (*MAD*) or *Qn*. Note that *MAD* may not suitably account for any potential skewness of the pairwise sequence patristic distance distribution as it inherently assumes symmetry about the median (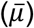). On the contrary, *Qn*, an alternative estimator of scale proposed by Rousseeuw & Croux (1993), is as robust as *MAD* (i.e. 50% breakdown point), calculated solely using the differences between the values in the distribution without needing a location estimate, and has been proven to be statistically more efficient in both Gaussian and non-Gaussian distributions relative to *MAD*.

### Integer linear programming model

Here, we fully elaborate the ILP model underlying PhyCLIP. Let *n*_1_, *n*_2_, …, *n*_*i*_, …, *n*_*N*_ be the set of binary variables indicating if subtree *i* satisfies the conditions for clustering as a clade (*n*_*i*_ = 1 if it does and *n*_*i*_ = 0 vice versa, Figure 2C). Each sequence *j* subtended by subtree *i* is also assigned a binary variable *j*_*j,i*_ indicating if the sequence is clustered under subtree *i* (*j*_*j,i*_ = 1 if *j* is clustered under node *i* and *j*_*j,i*_ = 0 vice versa, Figure 2C). PhyCLIP then formulates the phylogenetic clustering problem as an integer linear programming (ILP) model with the objective to maximise the number of sequences assigned with cluster membership:

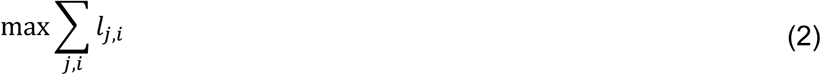

subject to the following constraints:

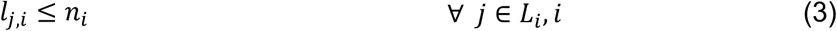

Constraint (3) stipulates that sequence *j* can be clustered under subtree *i* if and only if subtree *i* is a potential clade (*n*_*i*_ = 1).

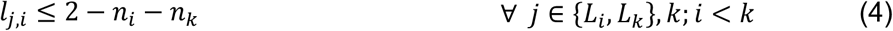

If sequence *j* is subtended by subtrees *i* and *k*, wherein *i* is ancestral to *k* and both nodes are potential clusters (*n*_*i*_ = *n*_X_ = 1), constraints (3) and (4) stipulate sequence *j* will not be clustered under the ancestor node *i*. Implementing these constraints across all pairwise combinations of subtrees subtending sequence *j* in turn constrains *j* to be clustered under the most descendant node *k* possible.

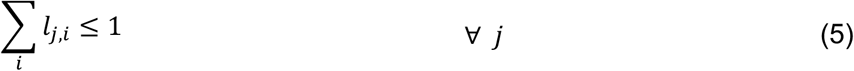

Constraint (5) stipulates that each sequence can only be clustered under a single subtree, hence abrogating any fuzzy clustering.

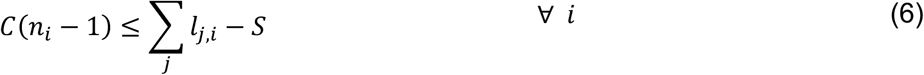

where *C* is any arbitrarily large positive constant. Constraint (6) requires all clusters to contain at least *S* number of taxa as defined by the user (Figures 1B and C).

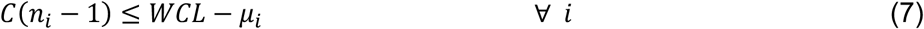

Constraint (7) ensures that *μ*_i_ of all clades fall below the stipulated *WCL* limit.

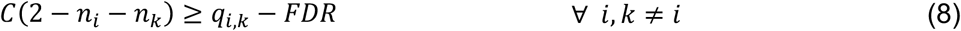

where *q*_*i,k*_ is the Benjamini-Hochberg corrected *p*-value testing if subtrees *i* and *k* are significantly divergent from one and another under the user-defined significance level, *FDR*. Constraint (8) is the inter-cluster divergence constraint. Inter-cluster divergence between subtrees *i* and *k* is tested under the null hypothesis that the pairwise sequence distance distributions of *i* and *k* are empirically equivalent to that if the two subtrees were clustered together. This can be done either by the putative Kolmogorov-Smirnov (KS) test or Kuiper’s test.

Although both tests are nonparametric, the Kuiper’s test statistic incorporates both the greatest positive and negative deviations between the two distributions whereas the KS test statistic is defined only by their maximum difference. As a result, the Kuiper’s test becomes equally sensitive to differences to the tails as well as the median of the distributions but the KS test works best when the distributions differ mostly at the median. In other words, the KS test is good at detecting *shifts* between the distributions but lacks the sensitivity to uncover *spreads* between the distributions characterised by changes in their tails. Kuiper’s test is, however, sensitive to detect both types of changes in distributions.

There are two scenarios under which *q*_*i,k*_ may be calculated:

i. Subtree *i* is ancestral to *k*. The hypothesis test assumes the null hypothesis that the pairwise sequence patristic distance distribution of subtree *k* is statistically identical to the pairwise sequence patristic distance distribution of its ancestor *i*.
ii. Neither subtree *i* nor *k* is an ancestor of the other. In this case, two hypothesis tests are carried out comparing the distribution of each subtree to the distribution of pairwise sequence patristic distance should both subtrees be combined as a single cluster and we take the more conservative *q_i,k_ = max{q_i,combined_, q_k,combined_*}.

### Nomenclature

Traversing the output clusters of PhyCLIP by pre-order of the input phylogeny, a unique number is assigned to any cluster with no immediate ancestral supercluster precursor to it (i.e. parent node of the cluster node is not part of any PhyCLIP clusters). Otherwise, the descendant cluster in question is designated as a *child cluster* should its membership size be >25^th^ percentile of PhyCLIP’s output cluster size distribution (i.e. for having proliferated in numbers substantial enough to be deemed a progeny cluster). Every child cluster of a supercluster is assigned a progeny number separated by a decimal point (e.g. 1.2 refers to the second child cluster of supercluster 1). On other hand, descendant clusters that fall below the cluster size cut-off are distinguished from child clusters as *nested clusters*, each assigned an address in the form of a parenthesized letter, alphabetised by tree traversal order, prefixed by its parent supercluster nomenclature (e.g. 1.1(c) refers to the third nested cluster of supercluster 1.1). Nested clusters in superclusters fundamentally have different properties from the sensitivity-induced nested clusters discussed in New Approach section and cannot be subsumed as it will violate the within-cluster limit of the parent supercluster. The structure of the resultant clustering topology is highlighted in Figure 3.

### Phylogenetic analyses

PhyCLIP’s performance was evaluated on an empirical dataset. The sequence datasets used to construct the haemagglutinin (HA) gene phylogenetic trees underlying the WHO/OIE/FAO nomenclature for the A/goose/Guangdong /1/1996 (Gs/GD/96)-like H5 avian influenza viruses were downloaded from GISAID (Anon 2008; WHO/OIE/FAO H5N1 Evolution Working Group 2012; WHO/OIE/FAO H5N1 Evolution Working Group 2014; Smith, Donis, and WHO/OIE/FAO H5 Evolution Working Group 2015). The primary analysis is based on the full dataset included in the 2009 (n=1224) and 2015 (n=4357) nomenclature updates. Viruses that were inconsistently included across WHO/OIE/FAO updates were followed up and included (WHO/OIE/FAO HN Evolution Working Gr 2009; Smith, Donis, andWHO/OIE/FAO H5 Evolution Working Group 2015). Sequences were curated based on criteria defined by the H5 nomenclature: sequences with more than 5 ambiguous nucleotides, with a sequence length shorter than 60% of the alignment, or with frameshifts or duplicated by name were removed. For the 2018 phylogeny, all avian and human viruses from the Gs/GD-like H5 lineage were downloaded from GISAID up to April 2018, including H5Nx subtypes H5N2, H5N3, H5N5, H5N6 and H5N8. An alternative filtering approach compared to the published WHO nomenclature approach was applied to ensure a dataset of high-quality sequences that would be robust to error in phylogenetic reconstruction as PhyCLIP is inherently sensitive to topological information. In this approach, duplicate sequences and sequences with a length below 95% of the full HA sequence or more than 1% ambiguous nucleotides were discarded. Sequences were aligned with MAFFT v7.397 and trimmed to the start of the mature protein (Katoh et al. 2002). Each sequence set was annotated with the WHO/OIE/FAOH5 nomenclature using LABEL(v0.5.2), and the version of the module corresponding to the nomenclature update of the dataset (e.g. H5v2015 module for the full tree from the nomenclature update in 2015) (Shepard et al. 2014). Maximum likelihood phylogenetic trees were constructed for each dataset with RAxML 8.2.12 under the GTR+GAMMA substitution model, and rooted to Gs/GD/96 (Stamatakis 2014). Phylogenetic trees were visualised using Figtree (http://tree.bio.ed.ac.uk/software/figtree/) and ggtree (Yu et al. 2017).

### Silhouette index

The silhouette index is based on the distance, here patristic distance, of each cluster member to other cluster members compared to the distance to its nearest neighbours (Rousseeuw 1987). Silhouette values approaching one indicate that the cluster member is correctly assigned, whereas values close to zero indicate that the sequence is equally matched to its neighbouring cluster. A negative Silhouette index indicates that the sequence is more closely related to the neighbouring cluster than to its fellow cluster members. Calculation of the silhouette index was performed in R (R Core Team 2016).

### Code availability

PhyCLIP is freely available on github (http://github.com/alvinxhan/PhyCLIP) and documentation can be found on the associated wiki page (http://github.com/alvinxhan/PhyCLIP/wiki).

## Supporting information

Figures S3, S5 and S11 in FigTree format

## Acknowledgments

We thank the GISAID Initiative and the influenza surveillance and research groups that openly shared the genetic sequence data that made this work possible (full acknowledgement table is available as supplementary). A.X.H. was supported by the A*STAR Graduate Scholarship programme from A*STAR to carry out his PhD work via collaboration between Bioinformatics Institute (A*STAR) and NUS Graduate School for Integrative Sciences and Engineering from the National University of Singapore. E.P. was funded by the Gates Cambridge Trust (Grant number OPP1144). S.M.S. was supported by the A*STAR HEIDI programme (Grant number: H1699f0013) and Bioinformatics Institute (A*STAR). C.A.R. was supported by University Research Fellowship from the Royal Society.

