## Supplementary Material for "Phylogenetic Clustering by Linear Integer Programming (PhyCLIP)"

### Mathematical primer on linear programming optimization and multiple optimal solutions

The linear programming model underlying PhyCLIP can be generalised as:

$$\max \sum_i c_i x_i \tag{i}$$

$$s.t. \quad \sum_i A_i x_i \leq b \tag{ii}$$

where *x* are variables, *b* is a constant, and *A* as well as *c* are coefficients. Intuitively, if $x \geq 0$ and $b \geq 0$, an optimal solution will always exist, which is the case for PhyCLIP.

We can rewrite (iii) and (iv) into equalities:

$$z - \sum_i c_i x_i = 0 \tag{iii}$$

$$\sum_i A_i x_i + s = b \tag{iv}$$

where $s$ is non-negative slack variable. As this point, we define *x* as a non-basic variable (appeared in >1 equation) and *s* as basic (appeared only in one equation). The basic solution is defined by setting all non-basic variables to zero. Hence, the initial basic solution is:

$$x_i = 0, s = b, z = 0$$

To increase *z*, one could increase the value of any variable $x_i$ so long as its corresponding coefficient in the objective equality is negative. This procedure to maximise *z* is known as the simplex algorithm.

In each iteration, a non-basic variable with a negative coefficient in the objective equality is selected as the entering variable (to become basic). For example, suppose $x_1$ is the entering variable, by Gaussian elimination:

$$z + \sum_{i=2} \left( \frac{c_1 A_i}{A_1} - c_i \right) x_i + \frac{c_1}{A_1} s = \frac{c_1 b}{A_1} \tag{v}$$

$$x_1 + \sum_{i=2} \frac{A_i}{A_1} x_i + \frac{s}{A_1} = \frac{b}{A_1} \tag{vi}$$

In this case, *s* is the leaving variable (to become non-basic). Setting all non-basic variables ($x_{i\geq 2}$ and $s$) as zero gives the following basic solution:

$$x_1 = \frac{b}{A_1}, x_{i\geq 2} = 0, s = 0, z = \frac{c_1 b}{A_1}$$

If all of the coefficients in the current objective equality (v) are non-negative (i.e. $\frac{c_1 A_{i\geq 2}}{A_1} - c_{i\geq 2} \geq 0$ and $\frac{c_1}{A_1} \geq 0$), the current basic solution is the optimal solution. Otherwise, the above procedure is repeated for another entering variable.

On the other hand, if a non-basic variable, say $x_m$, which coefficient in the objective equality happens to be zero upon converging to optimality, this implies $x_m$ can take alternate, non-negative values such that the optimal objective *z* remains unchanged. In other words, multiple optimal solutions exist. For example, suppose the following linear programming model:

$$\max x_1 + \frac{x_2}{3}$$

$$s.t. \quad 3x_1 + x_2 \leq 9$$

$$x_1 \geq 0, x_2 \geq 0$$

Intuitively, we know that there are multiple optimal solutions for this simple linear model. Formally, re-writing the above as equalities and selecting $x_1$ as the entering variable:

$$z + [\mathbf{0}]x_2 + \frac{s}{3} = 3$$

$$x_1 + \frac{x_2}{3} + \frac{s}{3} = 3$$

The basic solution becomes $x_1 = 3$, $x_2 = s = 0$ and $z = 3$. Since all coefficients in the objective equality are non-negative, this basic solution is also an optimal solution. However, as the coefficient of $x_2$ is equal to zero, $x_2$ can increase without changing the value of *z*. If we had selected $x_2$ as the entering variable instead:

$$z + [0]x_1 + \frac{s}{3} = 3 \quad \text{(vii)}$$

$$3x_1 + x_2 + s = 9 \quad \text{(viii)}$$

Now, the basic solution, which is also an alternate optimal solution, is $x_2 = 9$, $x_1 = s = 0$ and $z = 3$ remained unchanged.

### Supplementary Tables

**Table S1**: Optimal clustering result for the WHO/FAO/OIE 2015-update phylogeny, 2009-update phylogeny and H5Nx 2018 phylogeny respectively

**Table S2**: WHO/FAO/OIE clade designation for the 2015 Gs/GD-like H5N1 phylogeny

**Table S3**: Phylogenetic clusters designated by PhyCLIP present in each country represented in the WHO/FAO/OIE 2015 H5 nomenclature update

**Table S4**: Clustering properties of PhyloPart and ClusterPicker's optimal result

**Table S5**: Optimal clustering result for Phylopart on the 2009-update phylogeny

**Table S6**: GISAID acknowledgement table

### Supplementary Figures

**A**

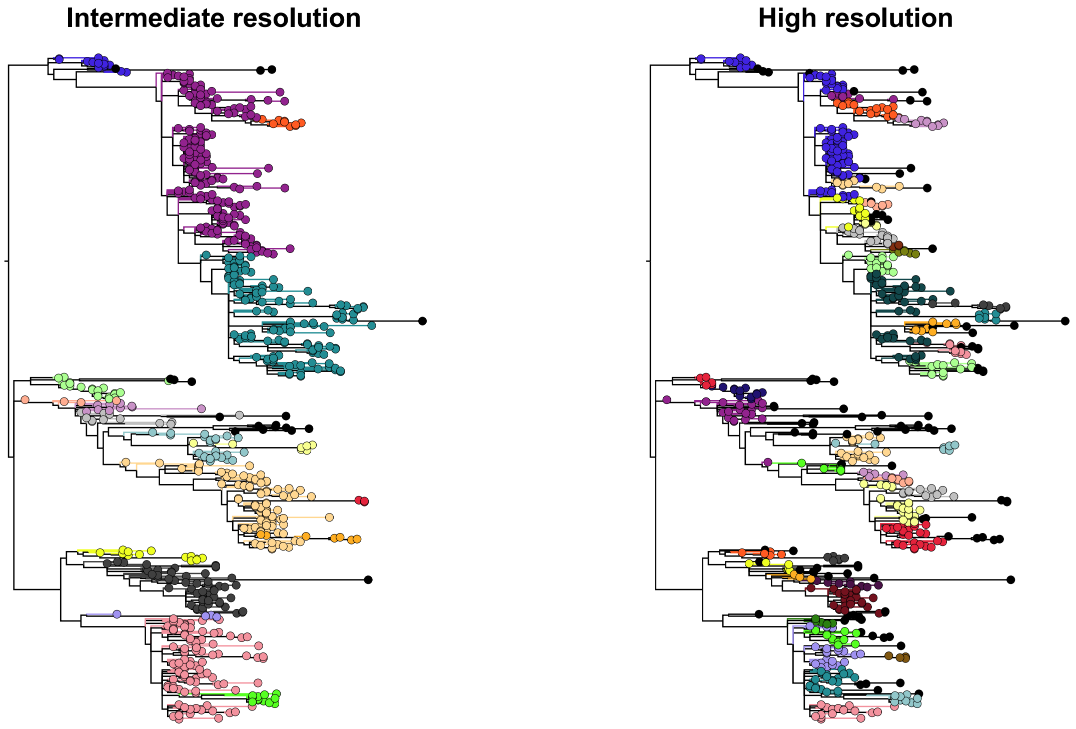


**B**

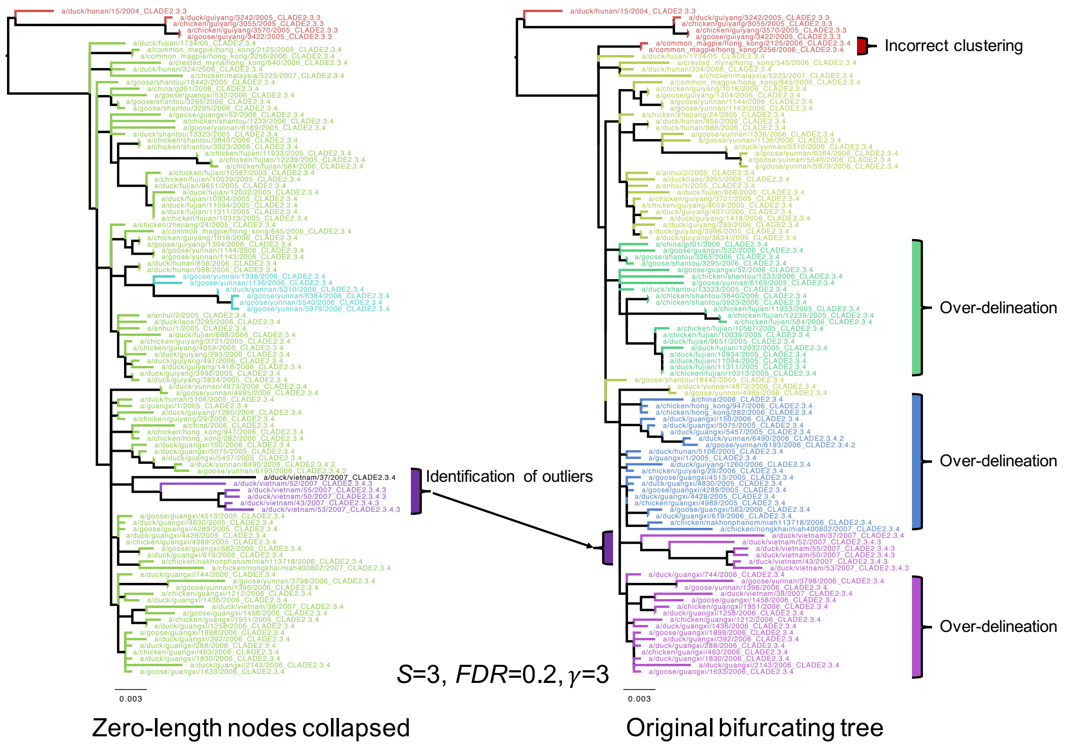


**Figure S1**: A. Example comparing between clustering results optimised for "intermediate" (more broadly defined clusters that are still well-separated but encompass the majority of data in the tree) and "high" (tree delineated into a large number of small, high confidence clusters with very low internal divergence) resolution by PhyCLIP. Distinct clusters are differentiated by different tip and branch colours (except for black branches/tips which represent unclustered taxa). B. Example of clustering results with and without collapsing non-terminal internal nodes with zero branch lengths, which are usually found in bifurcating trees representing polytomies. These zero-branch length internal nodes can lead to erroneous clustering and over-delineation in PhyCLIP's clustering results.

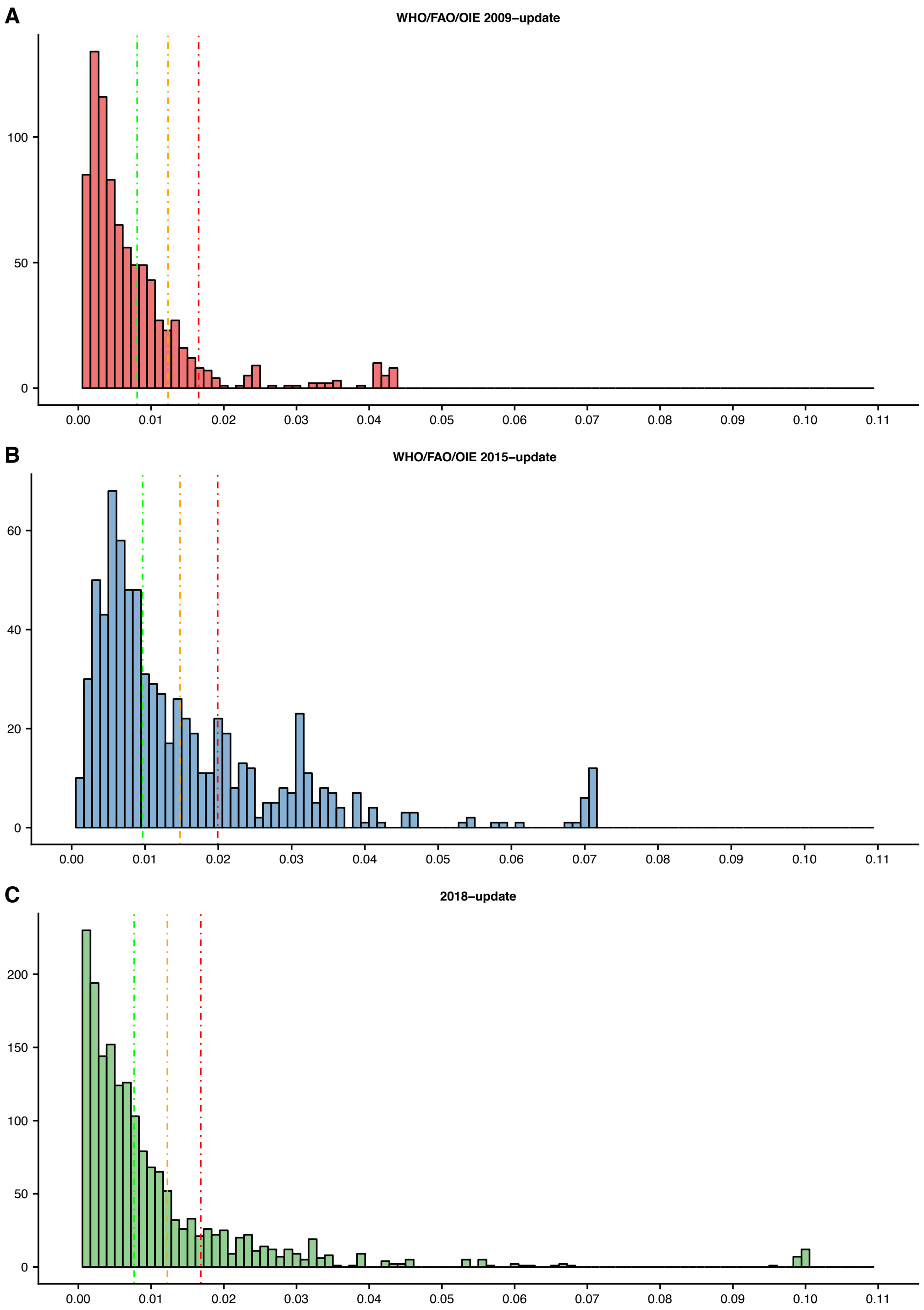


**Figure S2**: Distribution of the mean pairwise patristic distance of all of the internal nodes of the phylogeny above the minimum cluster criteria eligible for selection as putative clusters. A. WHO/FAO/OIE 2009-update of the H5 phylogeny. B. WHO/FAO/OIE 2015-update of the H5 phylogeny. C. 2018-update of the H5 phylogeny. The vertical lines designate the defined within-cluster limit at a gamma of one (green), two (orange) and three (red).

<See Figure_S3.tre.txt (FigTree File)>

**Figure S3**: PhyCLIP's optimal clustering result of the haemagglutinin phylogeny underlying the 2015 WHO/FAO/OIE nomenclature update of the Gs/GD-like H5N1 avian influenza viruses. Tips are coloured according to PhyCLIP's clustering designation. Tip names are appended with the WHO/FAO/OIE clade designation (_cladex) and PhyCLIP's cluster address (_clusterx).

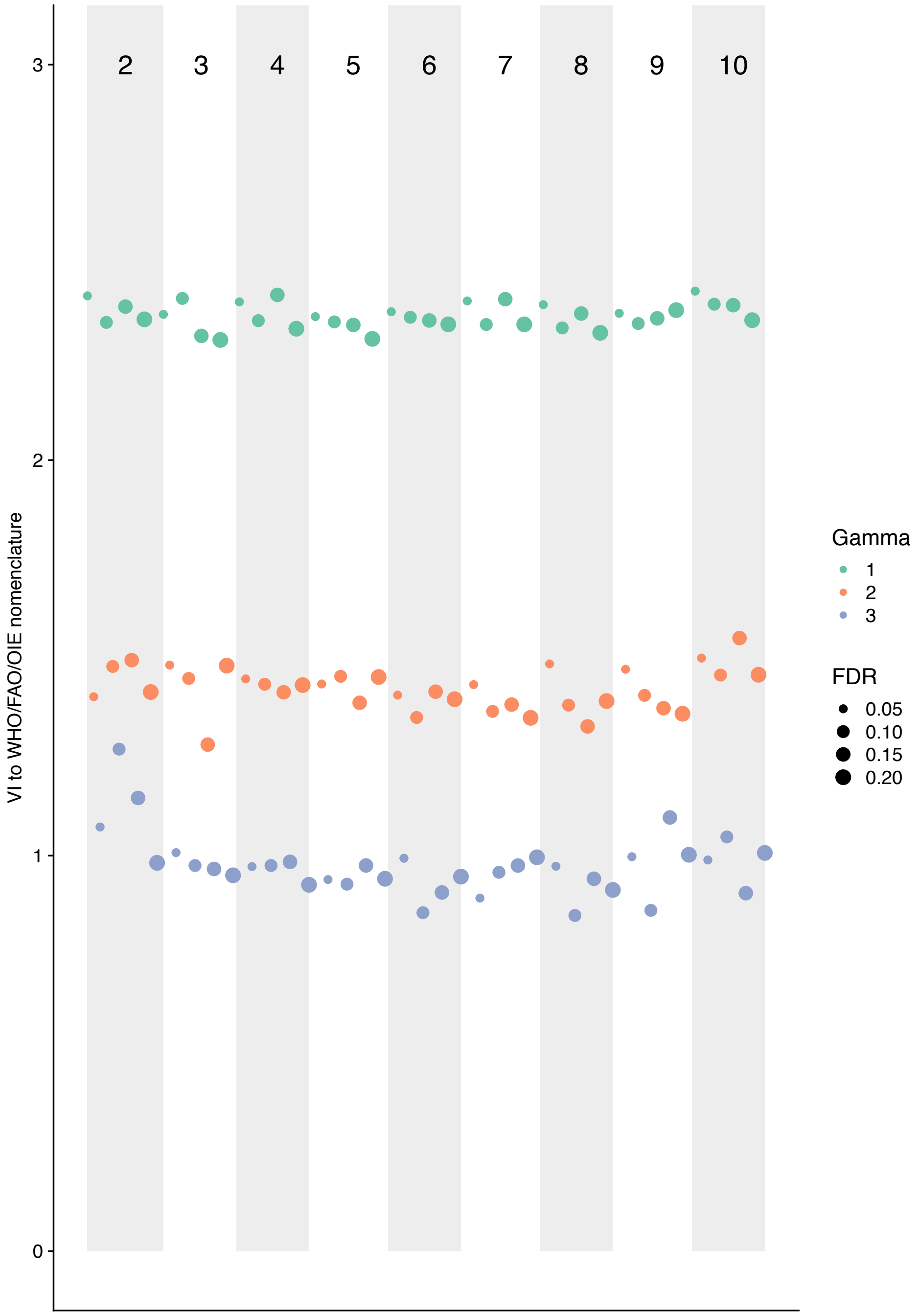


**Figure S4**: Comparison of PhyCLIP's clustering results to the WHO/FAO/OIE clade designation for the 2015 nomenclature update. The parameter set combinations ordered according to minimum cluster size, FDR and gamma are on the x axis. The banded background and x-axis superscript numbering indicate the minimum cluster size of the parameter set. Marker colour and size is indicative of the multiple of deviation and the false discovery rate respectively of the parameter set, as indicated by the legend.

<see Figure_S5.tre.txt (FigTree File)>

**Figure S5**: PhyCLIP's optimal clustering result of the haemagglutinin phylogeny underlying the 2009 WHO/FAO/OIE nomenclature update of the Gs/GD-like H5N1 avian influenza viruses. Tips are coloured according to PhyCLIP's clustering designation. Tip names are appended with the WHO/FAO/OIE clade designation (_cladex) and PhyCLIP's cluster address (_clusterx).

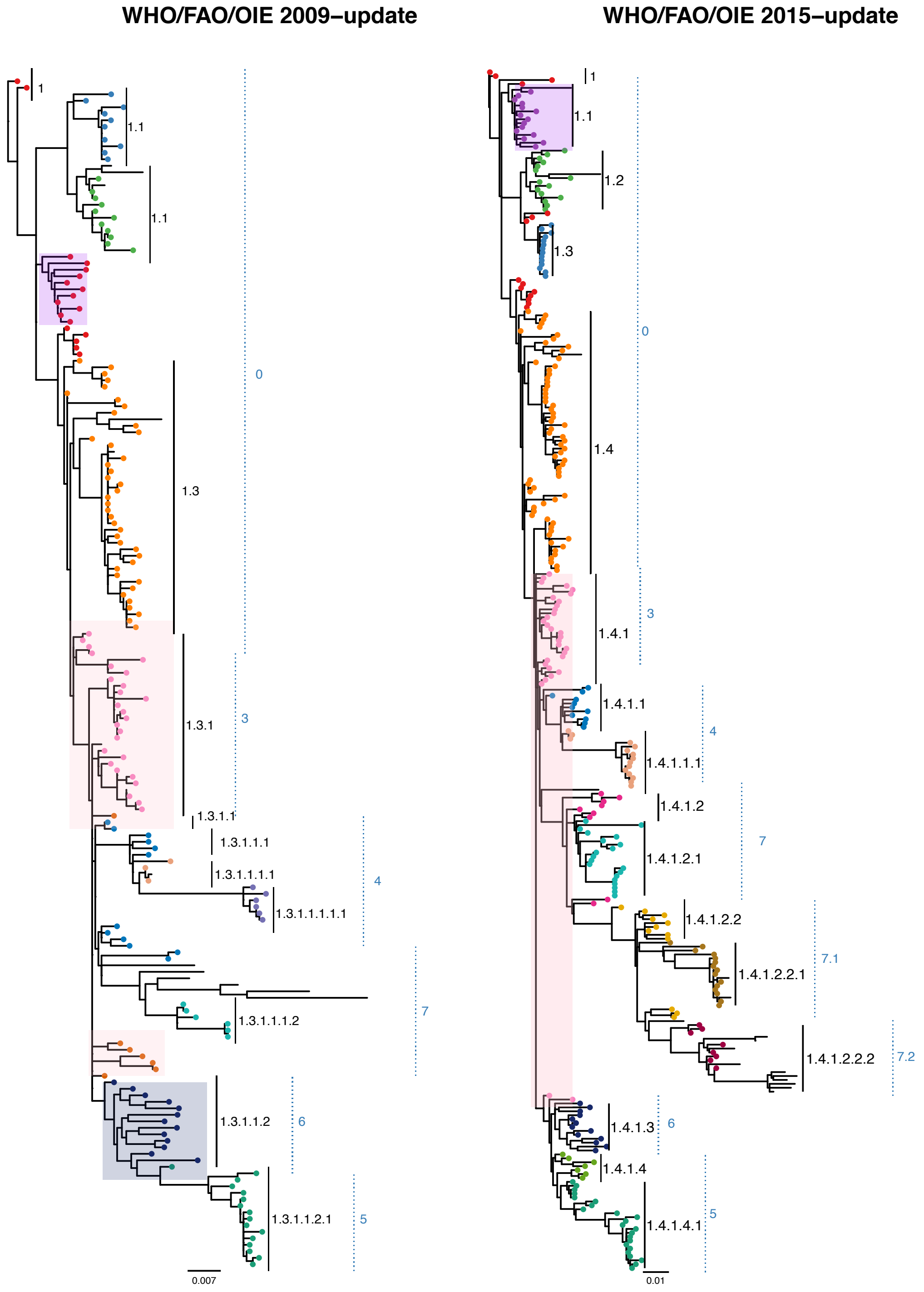


**Figure S6**: Comparison of changes between the optimal clustering result for the WHO/FAO/OIE 2015-update and 2009-update phylogenies owing to changes in the patristic distance distribution and topology for the clade 0, 3, 4, 5, 6, and 7 viruses. In the 2015 update phylogeny, descendent trunk viruses (indicated in pink) are incorporated into the supercluster 1.4.1 as their inclusion does not violate the within-cluster limit once the statistically distinct 1.4.1.1 and 1.4.1.2 and their descendants are dissociated. The 2009-update phylogeny defines a lower within-cluster limit, as the 2015-update phylogeny's distribution is shifted by the addition of newer, diverse viruses. In the 2009 phylogeny, these trunk viruses are basal to the clade highlighted in blue. They cannot be incorporated into cluster 1.3.1 (corresponding to 1.4.1) without violating the reduced within-cluster limit, and are considered as an independent cluster, cluster 1.3.1.1.2. This leads to shifts in the clustering inference drawn between 2009 and 2015. Topological difference between the trees can also underlie changes in clustering between the two phylogenies. PhyCLIP resolves an additional seed lineage, cluster 1.1, for the 2015-update phylogeny that forms part of the source population (cluster 1) in the 2009 phylogeny, as highlighted in purple. The orange tipped viruses highlighted in pink in the 2009 phylogeny form part of cluster 1.4.1 in the 2015 phylogeny, which underlie further topological changes between the trees that influences clustering inference. Clusters with membership consistent across the trees share a tip colour. Outliers are indicated as edges without tip-points.

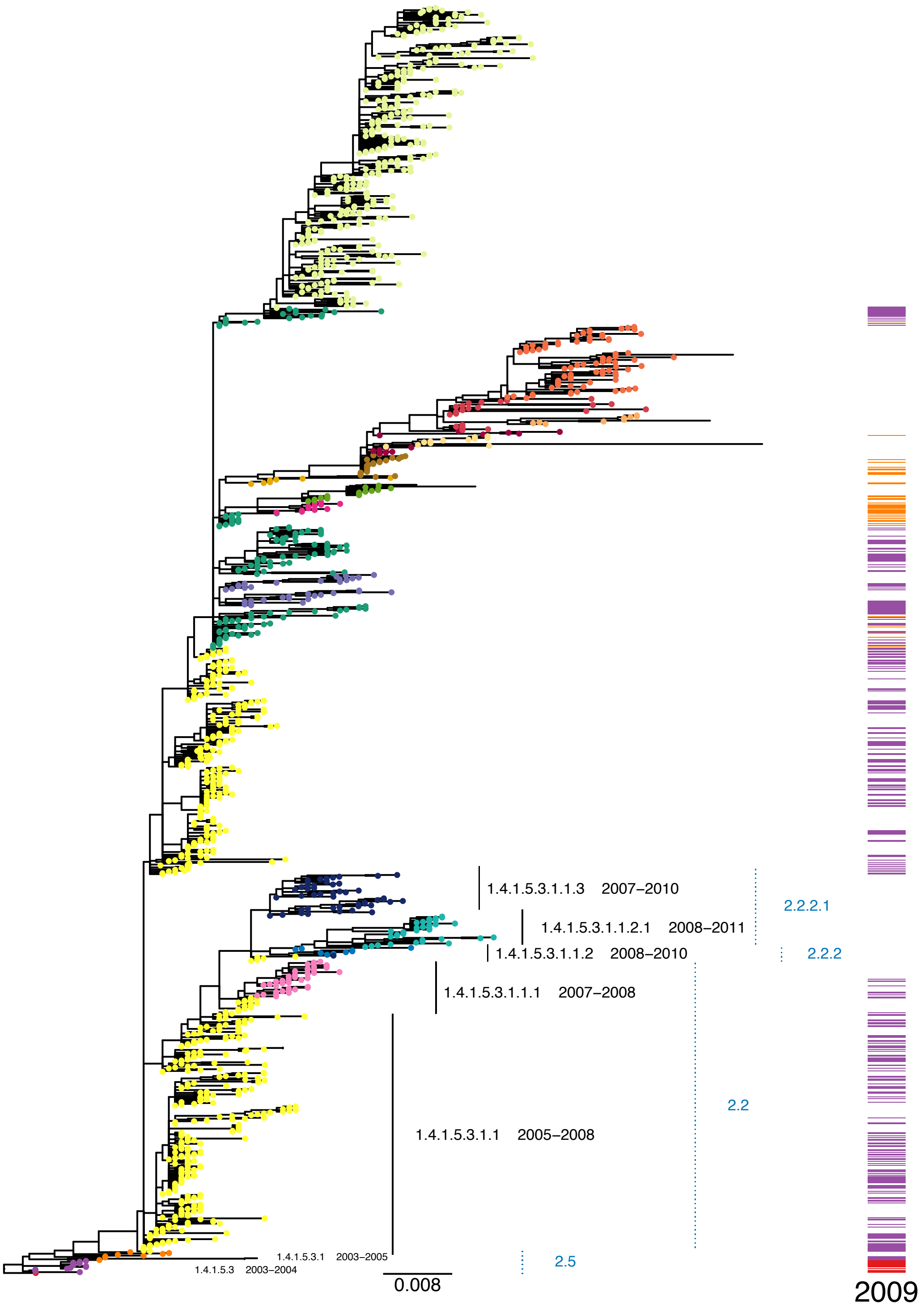


**Figure S7**: Comparison of changes between the optimal clustering result for the WHO/FAO/OIE 2015-update and 2009-update phylogenies owing to local addition of lineages for WHO for Clade 2.2x viruses. Tip colour indicates PhyCLIP's optimal cluster designation in the 2015-update phylogeny. The heat map indicates the PhyCLIP cluster designation for the viruses present in the 2015 also present in the 2009 phylogeny, based on the 2009 optimal clustering result. PhyCLIP only designates three clusters in the clade 2.2x viruses based on the 2009-update phylogeny, with one large cluster capturing most of the clade 2.2x viruses present in the 2009 phylogeny (indicated in blue in the heatmap). In the 2015 PhyCLIP cluster topology, this cluster is structured as a source population in a superclusters, cluster 1.4.1.5.3.1.1. The divergent descendants circulating post-2009 are dissociated to form descendant lineages e.g. cluster 1.4.1.5.3.1.1.1, 1.4.1.5.3.1.1.2 and 1.4.1.5.3.1.1.2.1, and 1.4.1.5.3.1.1.3. WHO/FAO/OIE clade designation is annotated in blue.

S8-A

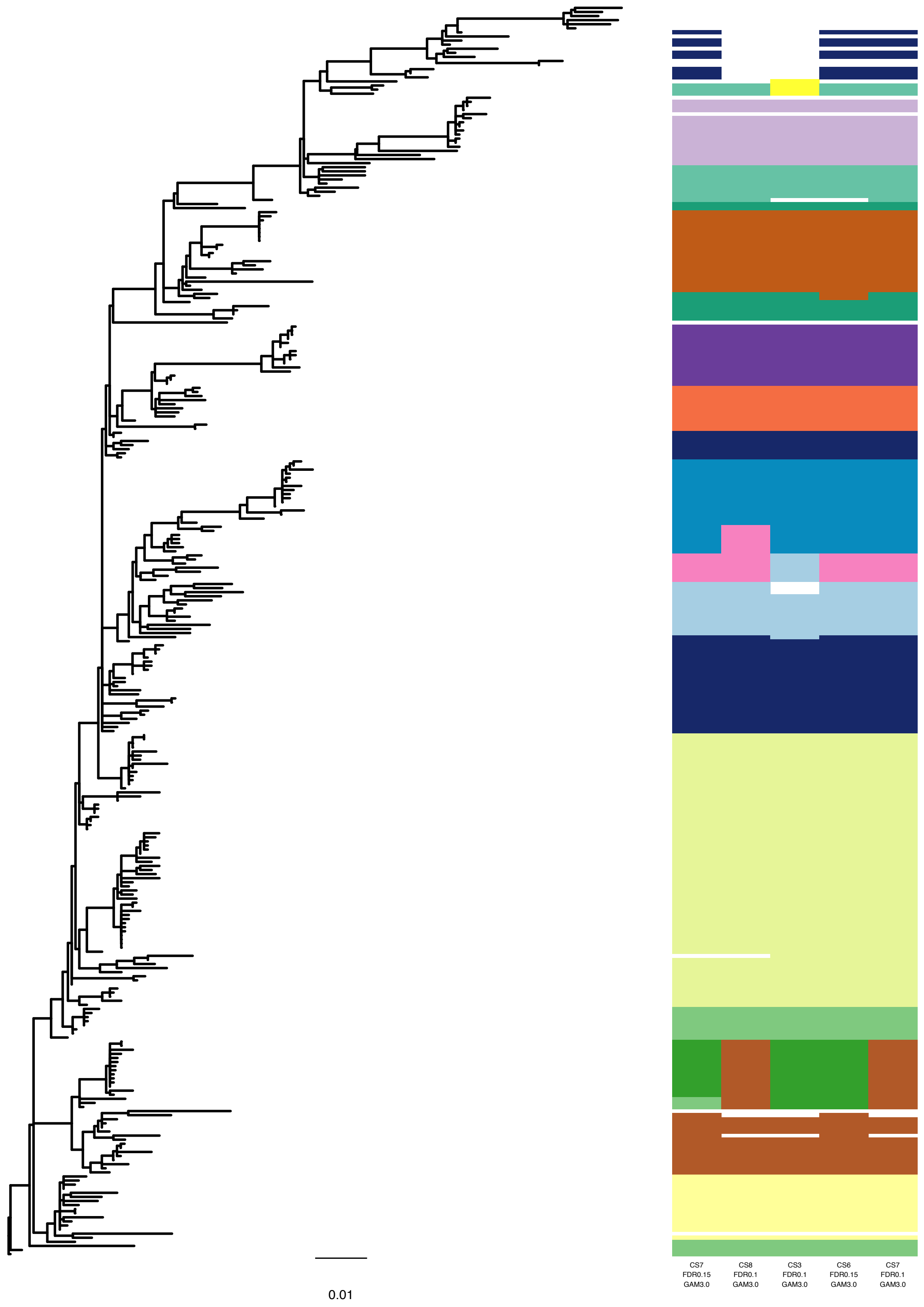
0.01
CS7
FDR0.15
GAM3.0
CS8
FDR0.1
GAM3.0
CS3
FDR0.1
GAM3.0
CS6
FDR0.15
GAM3.0
CS7
FDR0.1
GAM3.0

S8-B

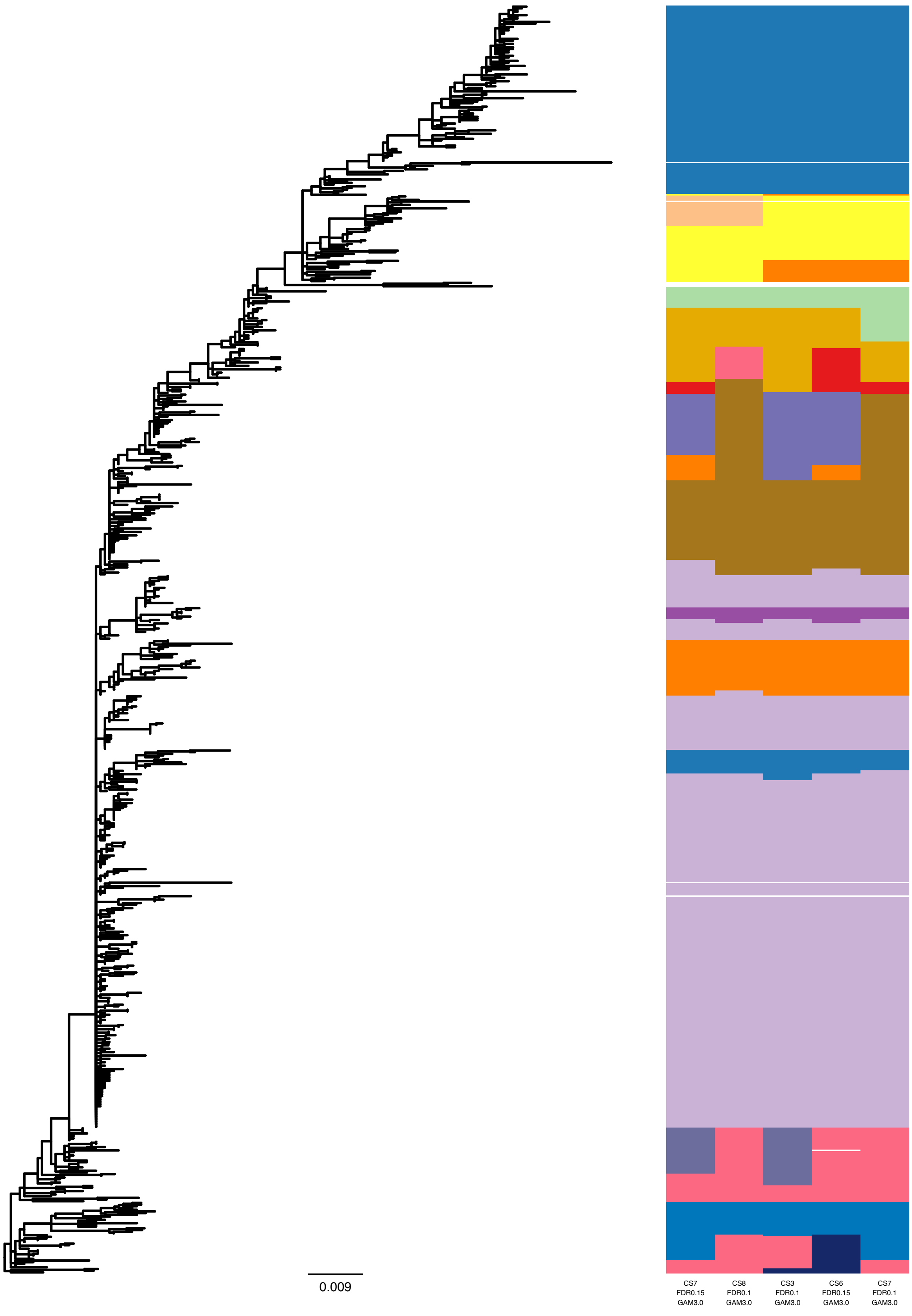
0.009
CS7
FDR0.15
GAM3.0
CS8
FDR0.1
GAM3.0
CS3
FDR0.1
GAM3.0
CS6
FDR0.15
GAM3.0
CS7
FDR0.1
GAM3.0

**S8-C**

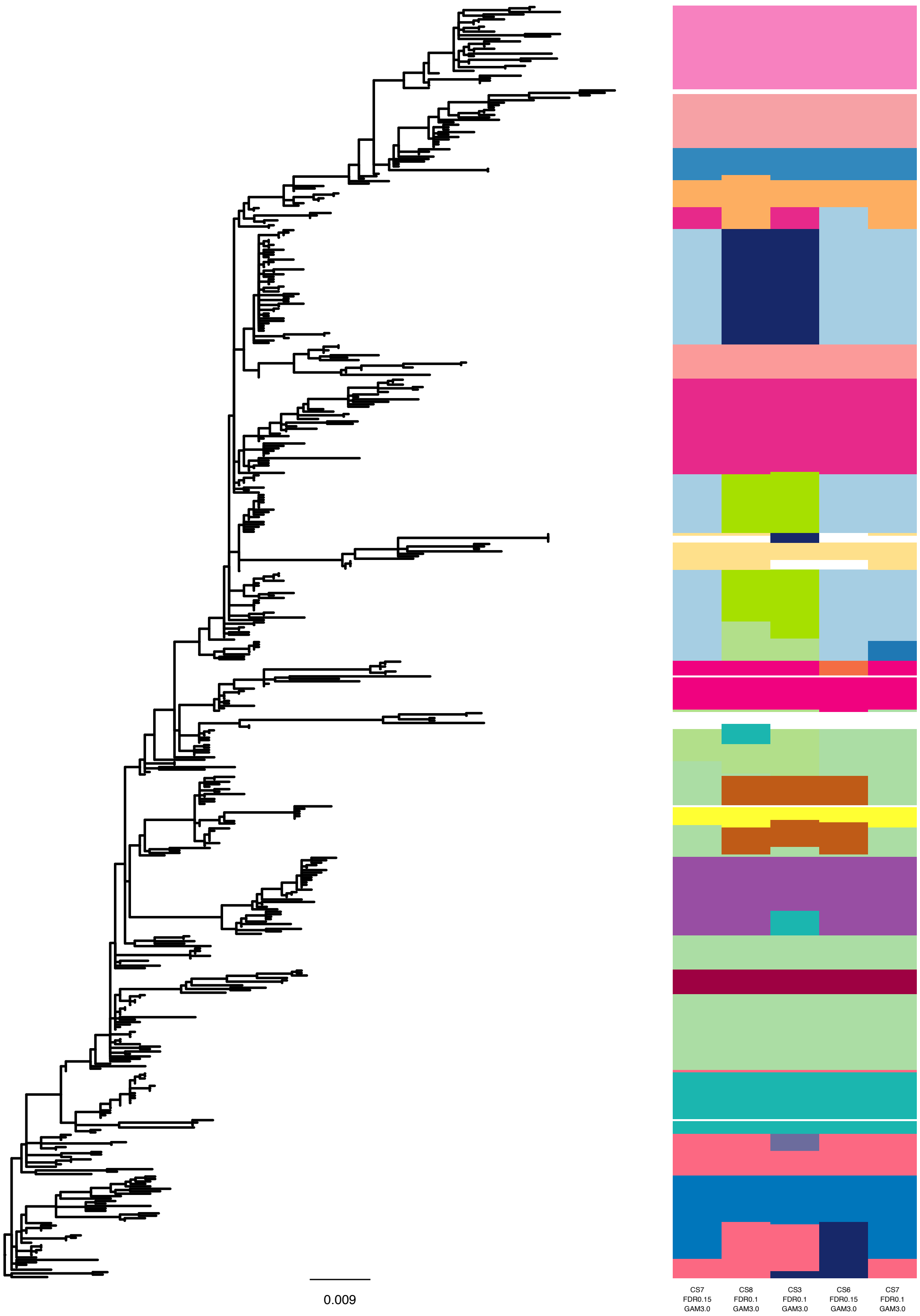
0.009
CS7
FDR0.15
GAM3.0
CS8
FDR0.1
GAM3.0
CS3
FDR0.1
GAM3.0
CS6
FDR0.15
GAM3.0
CS7
FDR0.1
GAM3.0

S8-D

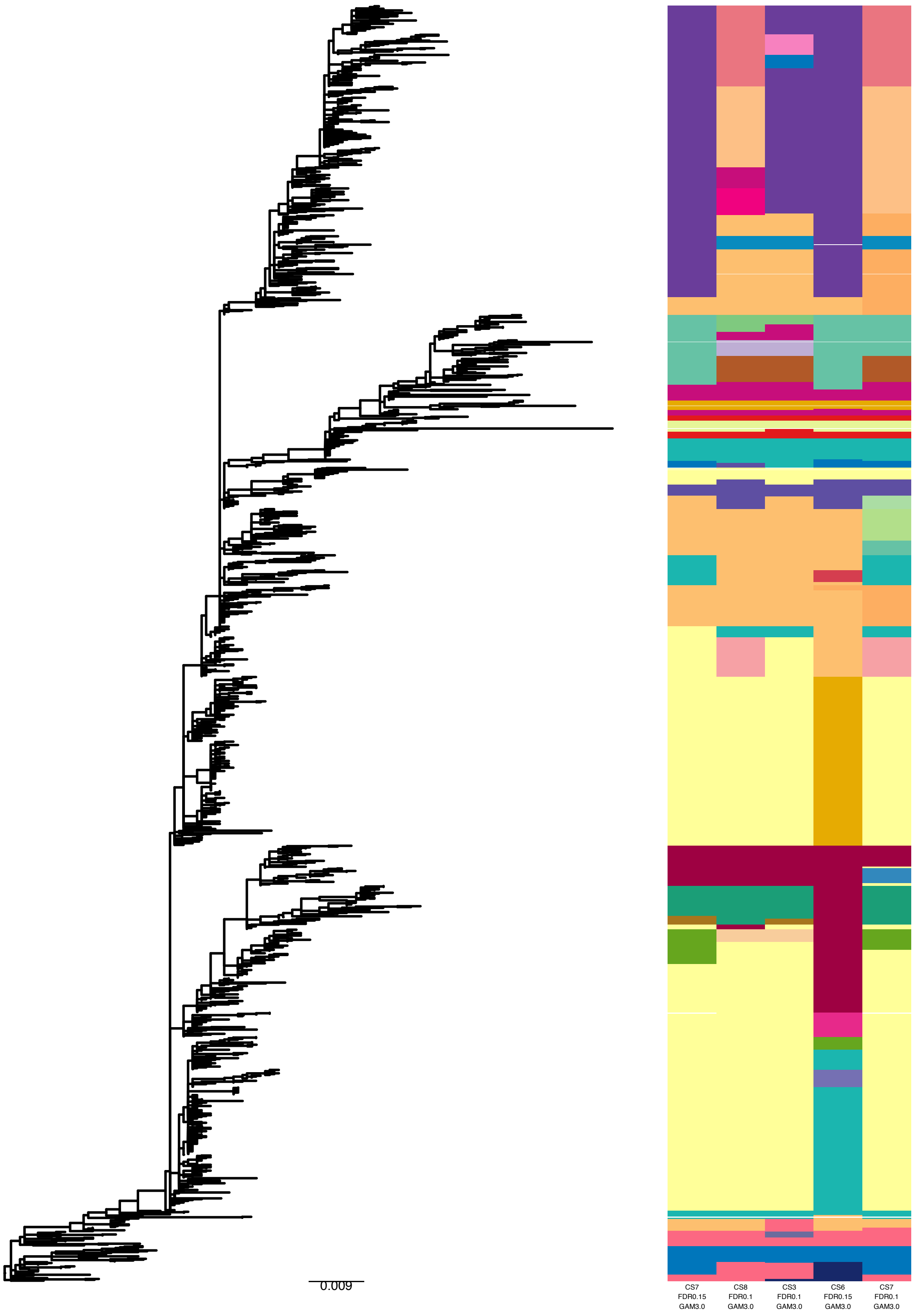
0.009
CS7
FDR0.15
GAM3.0
CS8
FDR0.1
GAM3.0
CS3
FDR0.1
GAM3.0
CS6
FDR0.15
GAM3.0
CS7
FDR0.1
GAM3.0

**S8-E**

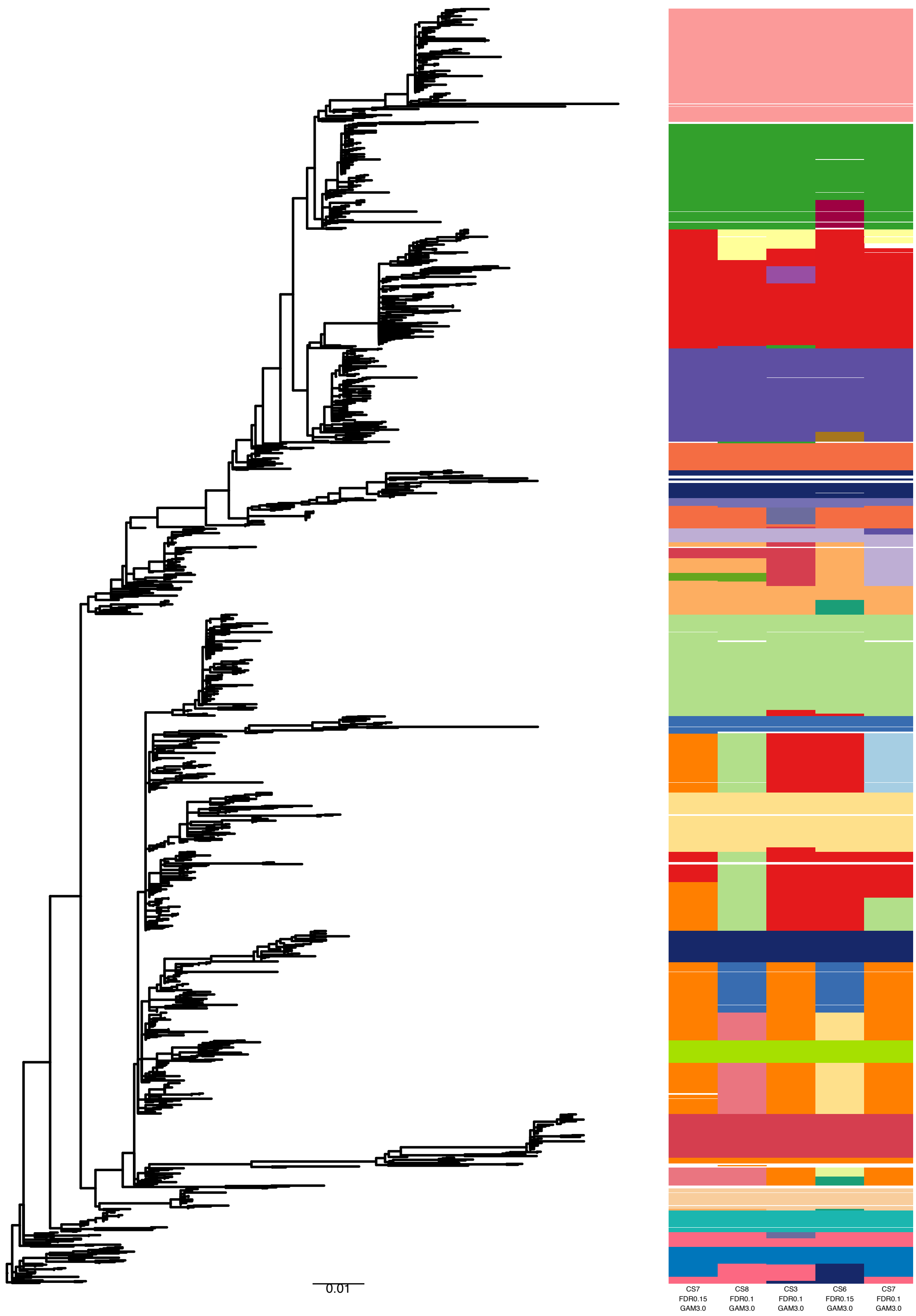


**Figure S8**: Comparison of optimal (*S*=7, *FDR*=0.15, $\gamma$=3) and top four sub-optimal (in order of sub-optimality – *S*=8, *FDR*=0.10, $\gamma$=3; *S*=3, *FDR*=0.10, $\gamma$=3; *S*=6, *FDR*=0.15, $\gamma$=3; *S*=6, *FDR*=0.10, $\gamma$=3) clustering results of the WHO/FAO/OIE 2015-update H5 phylogeny. The heat map indicates PhyCLIP cluster designation. Colours in the sub-optimal results are matched to the corresponding cluster designation found in the optimal result (i.e. largest possible cluster with >50% matched sequences). A unique colour is given If no matching cluster is found in the optimal result. (a) Classical clade viruses (Clades 0, 3, 4, 5, 6 and 7.x). (b) Second supercluster leading to Clade 1 viruses. (c) Second supercluster leading to Clade 2.1.x viruses. (d) Second supercluster leading to Clade 2.2.x viruses. (e) Second supercluster leading to Clade 2.3.x viruses.

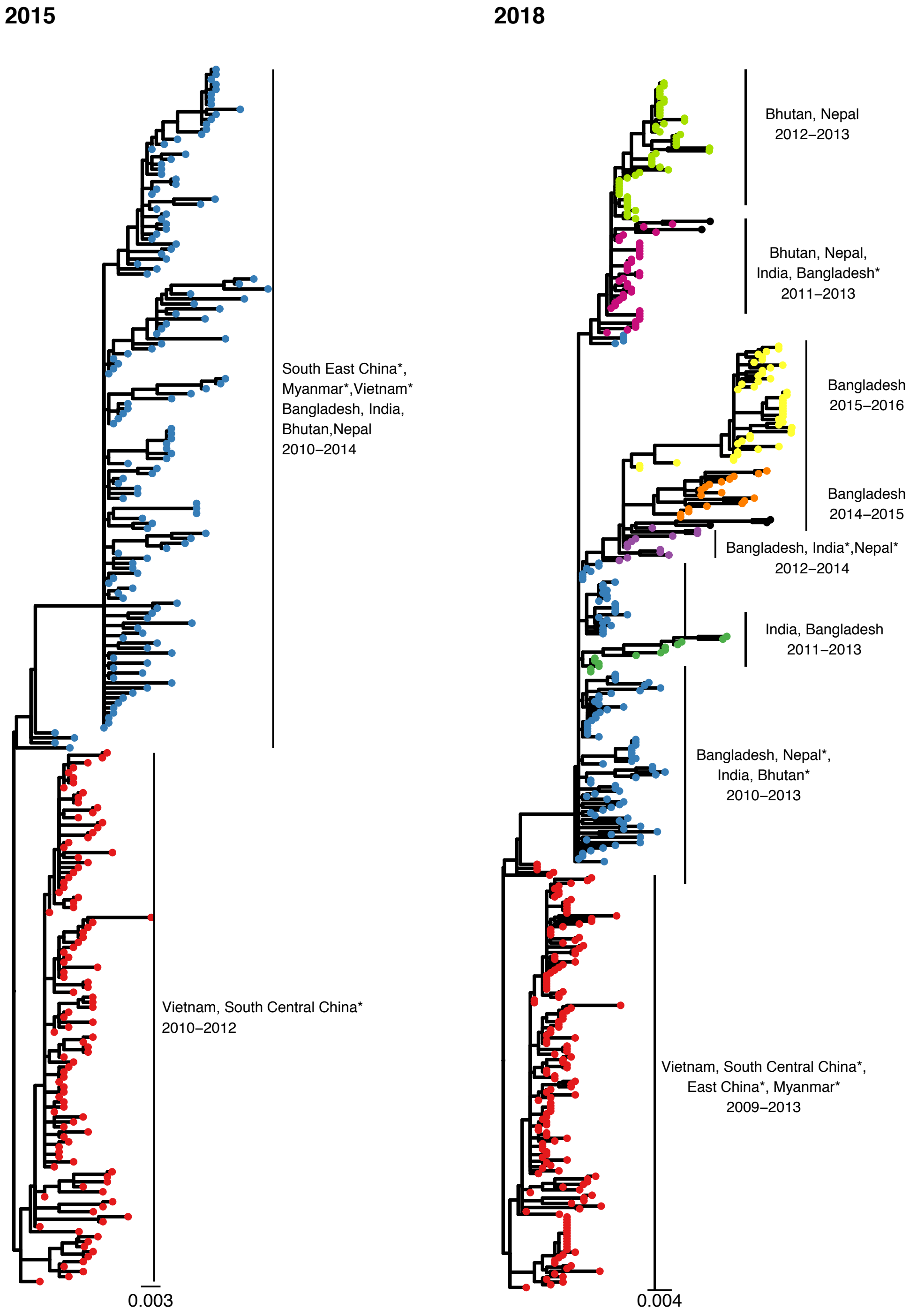


**Figures S9**: PhyCLIP's delineation of WHO/FAO/OIE demarcated clade 2.3.2.1a in the WHO/FAO/OIE 2015-update phylogeny and 2018 H5Nx phylogeny. Tips are coloured according to PhyCLIP's cluster designation. Corresponding clusters in 2015 and 2018 are matched in colour. The tips coloured in black are designated outliers. Countries represented by single viruses in the cluster are indicated with an asterisk.

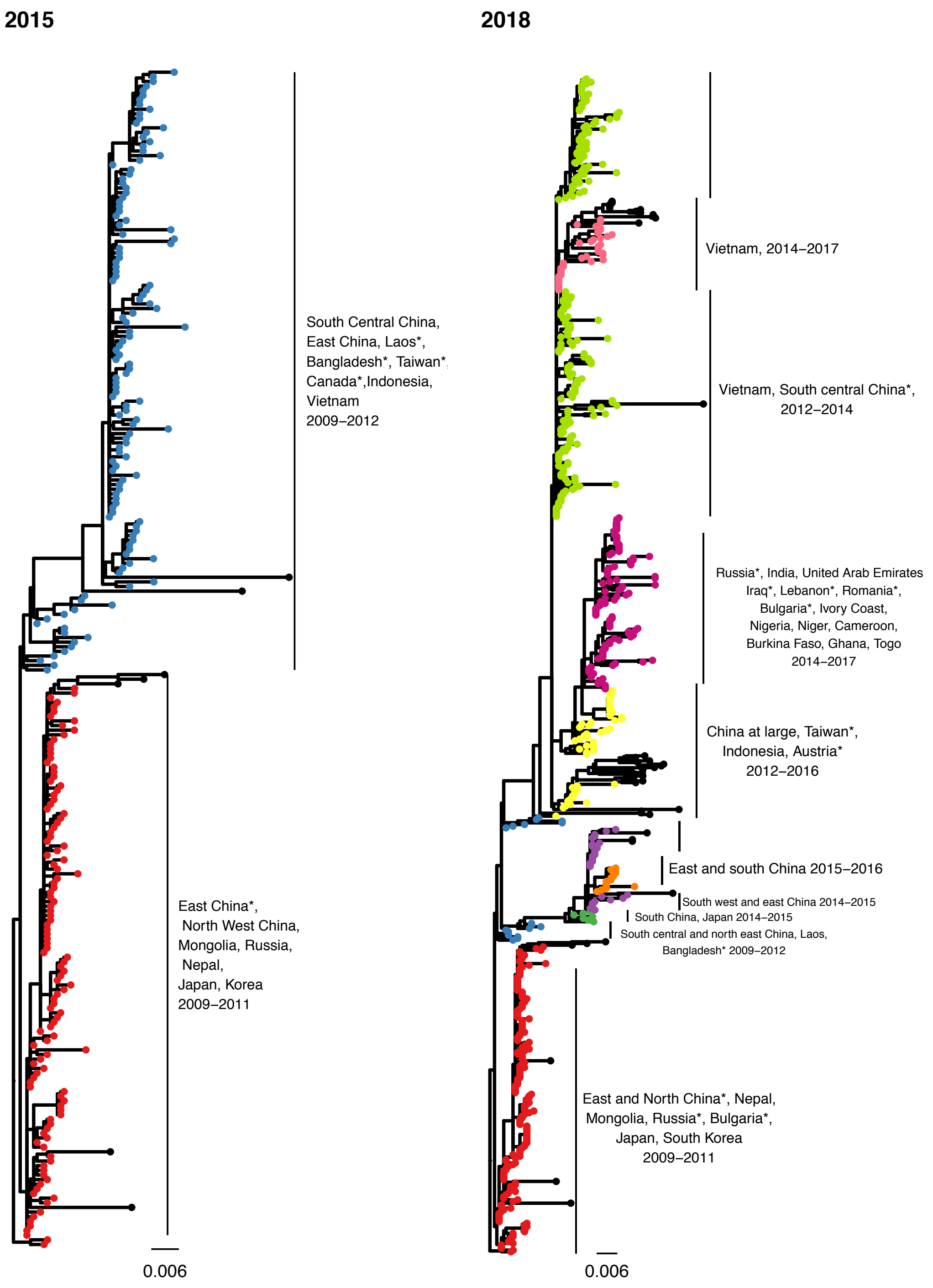


**Figures S10**: PhyCLIP's delineation of WHO/FAO/OIE demarcated clade 2.3.2.1c (Figure S10) in the WHO/FAO/OIE 2015-update phylogeny and 2018 H5Nx phylogeny. Tips are coloured according to PhyCLIP's cluster designation. Corresponding clusters in 2015 and 2018 are matched in colour. The tips coloured in black are designated outliers. Countries represented by single viruses in the cluster are indicated with an asterisk.

<see Figure_S11.tre.txt (FigTree File)>

**Figure S11**: Phyclip's optimal clustering result of the 2018 updated haemagglutinin phylogeny of the Gs/GD-like H5Nx avian influenza viruses. Tips are coloured according to PhyCLIP's clustering designation. Tip names are appended with the WHO/FAO/OIE clade designation (_cladex) and PhyCLIP's cluster address (_clusterx).

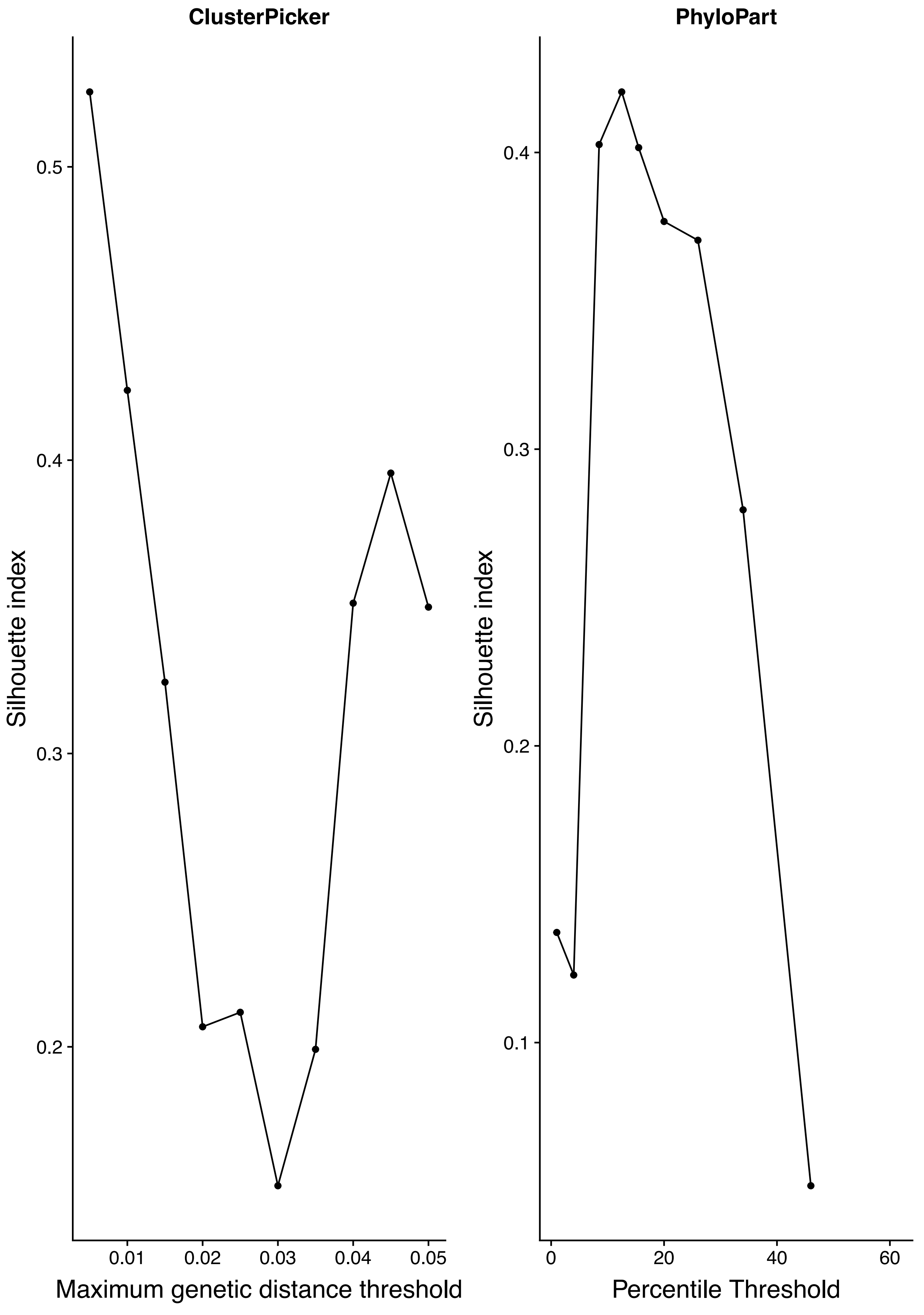


**Figure S12:** Silhouette index for ClusterPicker and PhyloPart cluster partitions designated over a range of distance thresholds. ClusterPicker's distance threshold is measured in maximum genetic distance, with the silhouette index maximised at 0.005. PhyloPart's distance threshold is defined as a percentile of the global pairwise patristic distance distribution, and is maximised at 12.5%, representing a median patristic distance threshold of 0.025.

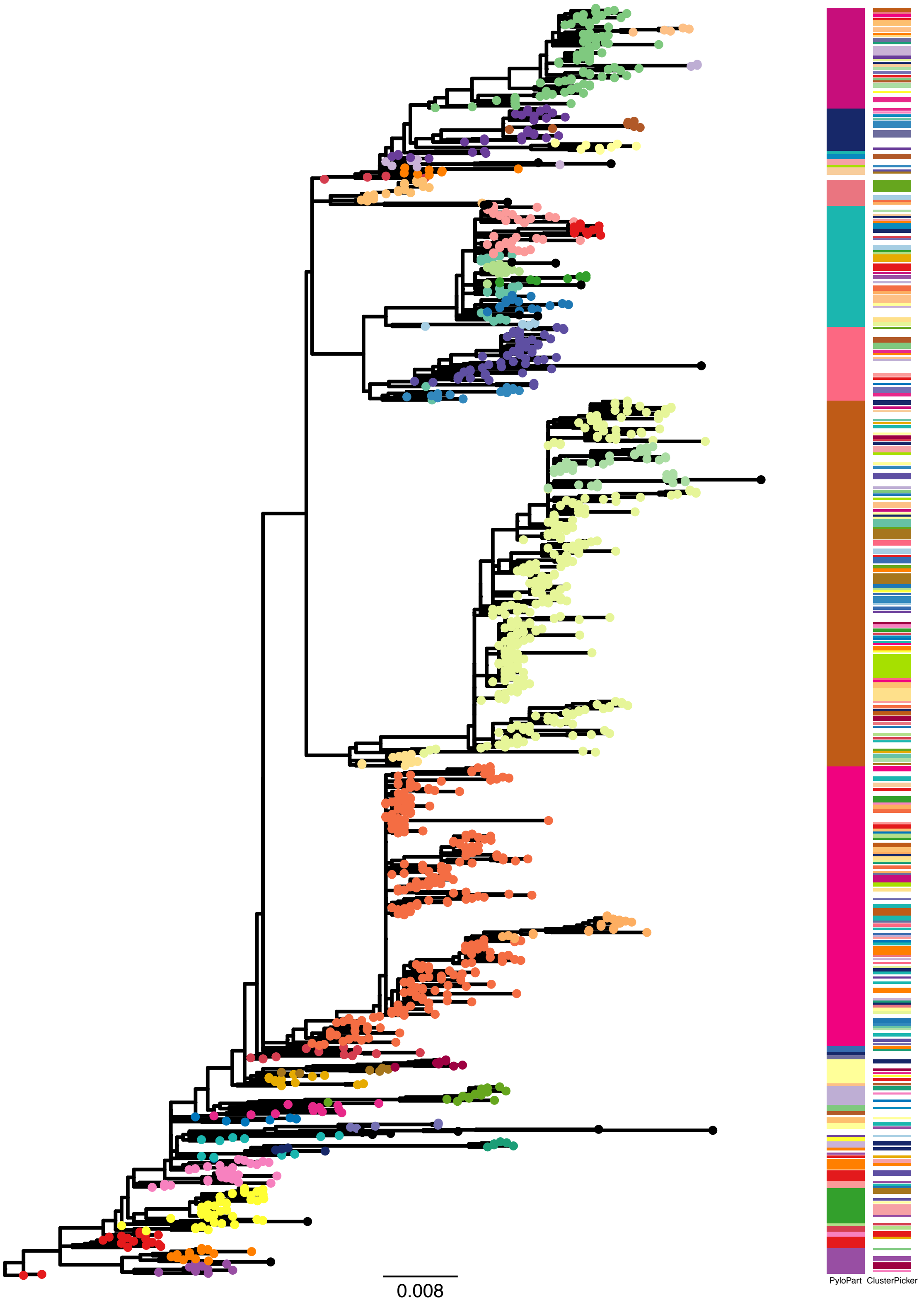


**Figure S13**: Comparison of PhyCLIP's optimal clustering result to PhyloPart and ClusterPicker's optimal clustering results, obtained by maximisation of the clustering partition silhouette index. The comparison was performed on the phylogeny underlying the 2009 WHO/FAO/OIE H5 nomenclature update. Tree tips are coloured according to PhyCLIP's cluster designation. Outliers are indicated with black tips. The heatmap indicates cluster designation according to PhyloPart, with a median patristic distance threshold of 0.025, (left) and ClusterPicker, with a maximum genetic distance of 0.005 subs/site (right).
